# Active Efficient Coding Explains the Development of Binocular Vision and its Failure in Amblyopia

**DOI:** 10.1101/571802

**Authors:** Samuel Eckmann, Lukas Klimmasch, Bertram E. Shi, Jochen Triesch

**Affiliations:** Frankfurt Institute for Advanced Studies, Frankfurt am Main, Germany; Department of Computer Science and Mathematics, Goethe University, Frankfurt am Main, Germany; Max Planck Institute for Brain Research, Frankfurt am Main, Germany; Department of Electronic and Computer Engineering, Hong Kong University of Science and Technology, Clear Water Bay, Hong Kong

**Keywords:** efficient coding, active perception, amblyopia, vergence, accommodation

## Abstract

The development of vision during the first months of life is an active process that comprises the learning of appropriate neural representations and the learning of accurate eye movements. While it has long been suspected that the two learning processes are coupled, there is still no widely accepted theoretical framework describing this joint development. Here we propose a computational model of the development of active binocular vision to fill this gap. The model is based on a new formulation of the *Active Efficient Coding* theory, which proposes that eye movements, as well as stimulus encoding, are jointly adapted to maximize the overall coding efficiency. Under healthy conditions, the model self-calibrates to perform accurate vergence and accommodation eye movements. It exploits disparity cues to deduce the direction of defocus, which leads to co-ordinated vergence and accommodation responses. In a simulated anisometropic case, where the refraction power of the two eyes differs, an amblyopia-like state develops, in which the foveal region of one eye is suppressed due to inputs from the other eye. After correcting for refractive errors, the model can only reach healthy performance levels if receptive fields are still plastic, in line with findings on a critical period for binocular vision development. Overall, our model offers a unifying conceptual framework for understanding the development of binocular vision.

**Significance Statement:** Brains must operate in an energy-efficient manner. The efficient coding hypothesis states that sensory systems achieve this by adapting neural representations to the statistics of sensory input signals. Importantly, however, these statistics are shaped by the organism’s behavior and how it samples information from the environment. Therefore, optimal performance requires jointly optimizing neural representations and behavior, a theory called *Active Efficient Coding*. Here we test the plausibility of this theory by proposing a computational model of the development of binocular vision. The model explains the development of accurate binocular vision under healthy conditions. In the case of refractive errors, however, the model develops an amblyopia-like state and suggests conditions for successful treatment.

---

Our brains are responsible for 20% of our energy consumption (1). Therefore, organizing neural circuits to be energy efficient may provide a substantial evolutionary advantage. One means of increasing energy efficiency in sensory systems is to attune neural representations to the statistics of sensory signals. Based on this *Efficient Coding Hypothesis* (2), numerous experimental observations in different sensory modalities have been explained (3, 4). For instance, it has been shown that receptive field properties in the early visual pathway can be explained through models that learn to efficiently encode natural images (5, 6). These findings have extended classic results showing that receptive field shapes in visual cortex are highly malleable and a product of the organism’s sensory experience (7–10).

Importantly, however, animals can shape the statistics of their sensory inputs through their behavior (Fig. 1). This gives them additional degrees of freedom to optimize coding efficiency by jointly adapting their neural representations and behavior. This idea has recently been advanced as *Active Efficient Coding* (11, 12). It can be understood as a generalization of the efficient coding hypothesis (2) to active perception (13). Along these lines, *Active Efficient Coding* models have been able to explain the development of visual receptive fields and the self-calibration of smooth-pursuit and vergence eye movements (11, 12). This has been achieved by optimizing the neural representation of the sensory signal statistics, while simultaneously, via eye movements, optimizing the statistics of sensory signals themselves, for maximal coding efficiency.

**Fig. 1.**
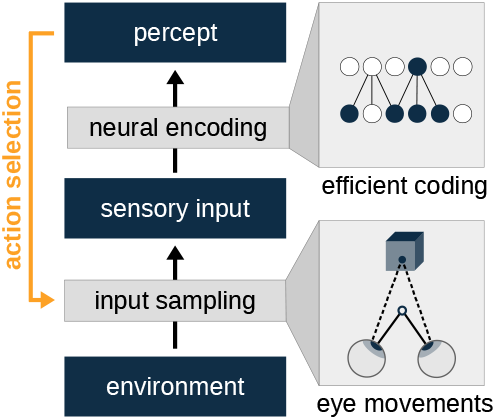
The action-perception loop in *Active Efficient Coding*. The sensory input is obtained by sampling input signals from the environment, e.g., via eye movements. A percept is formed by neural encoding which drives the selection of actions and thereby shapes the sampling process. Therefore, perception depends on both, neural encoding and active input sampling. Classic efficient coding theories do not consider the active sampling component (orange).

In our formulation of *Active Efficient Coding*, we maximize coding efficiency as measured by the Shannon mutual information *I*(*R, C*) between the retinal stimulus represented by retinal ganglion cell activity *R* and its cortical representation *C* under a limited resource constraint. The mutual information can be decomposed as:

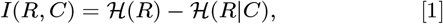

where 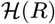 is the entropy of the retinal response and 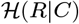 its conditional entropy given the cortical representation.

Prior formulations focused on minimizing the conditional entropy 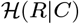 only (6). 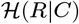 is a measure of the information that is lost, i.e., not represented in the cortical encoding. The limitation of this prior formulation is that this quantity can be minimized by simply reducing 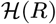, the entropy of the retinal response, since 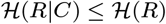. Thus, an active agent could minimize 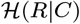 by, e.g., defocusing or closing the eyes altogether. In the free-energy and predictive processing literature, this is known as the “dark room problem” (14, 15). In our formulation, maximizing *I*(*R, C*) is achieved by maximizing 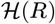 and minimizing 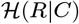 simultaneously, thus avoiding this problem. We demonstrate this approach through a concrete model of the development of binocular vision including the simultaneous calibration of vergence and accommodation control.

Indeed, newborns have difficulties bringing objects into focus and cannot yet verge their eyes properly (16). How infants manage to self-calibrate their control mechanisms while interacting with their visual environment is currently unknown. Additionally, in certain medical conditions, the calibration of vergence and accommodation control is impaired. For example, anisometropia describes a difference in the refractive error between the eyes. If not corrected early during development, this can evoke amblyopia: a disorder of the developing visual system that is characterized by an interocular difference in visual acuity that is not immediately resolved by refractive correction. Amblyopia can be associated with a loss of stereopsis and in severe cases leads to monocular blindness (17). Furthermore, vergence and accommodation eye movements are either less accurate or completely absent (18, 19).

Although there have been recent advances in the treatment of amblyopia (20, 21), existing treatment methods do not lead to satisfactory outcomes in all patients. This is aggravated by the fact that treatment success strongly depends on the stage of neural circuit maturation (20). When young patients are still in a critical period of visual cortex plasticity (10), they often recover after refractive errors are corrected, while adults mostly remain impeded (22, 23).

The above findings are all readily explained by our model. Under healthy conditions, our model develops accurate vergence and accommodation eye movements. When the model is impaired due to strong monocular hyperopia, we observe that an amblyopia-like state develops. We show that this is due to the abnormal development of binocular receptive fields in the model and demonstrate that healthy binocular vision is regained as the receptive fields readapt after refraction correction. However, if the sensory encoding is no longer plastic and does not adapt to the changes in the visual input statistics, suppression prevails. Overall, our model suggests that coding efficiency may provide a unifying explanation for the development of binocular vision.

## Model Formulation

The *Active Efficient Coding* model we propose has a modular structure (Fig. 2). A cortical coding module models the learning of an efficient representation *C* of the binocular retinal representation *R* by minimizing the conditional entropy 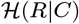. At the same time, an accommodation reinforcement learning module maximizes 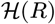 and a vergence reinforcement learning module minimizes 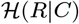 (see Eq. 1). All three modules are plastic and adjust simultaneously in response to changes in the sensory input statistics. The exact choice of the algorithms is not important for the model to function. In fact, different cortical coding and reinforcement learning models have been successfully applied in previous *Active Efficient Coding* models (24, 25).

**Fig. 2.**
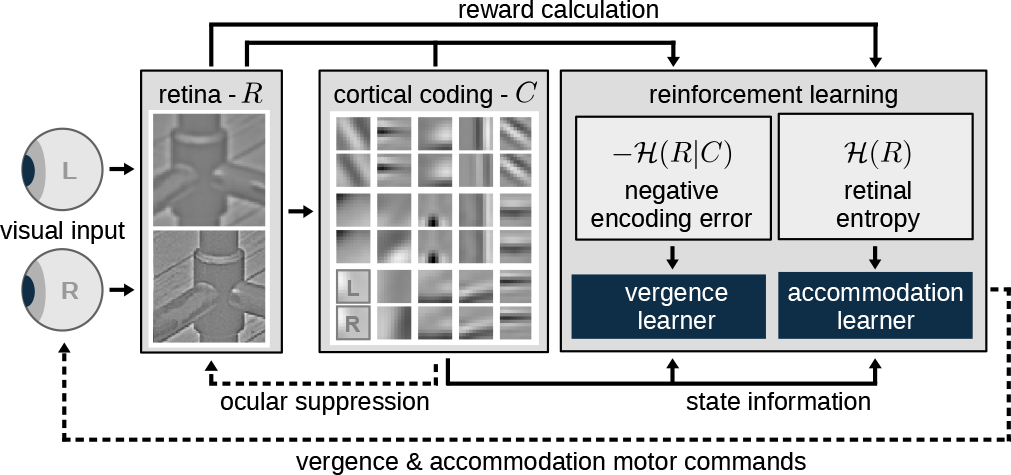
Model architecture with solid arrows representing the flow of sensory information and dashed arrows representing the flow of control commands. Sampled input images with given defocus blur and disparity are whitened at the retinal stage *R* and contrast adjusted through an interocular suppression mechanism based on the recent history of cortical activity (left). Thereafter, they are encoded by a set of binocular neurons which represents the cortical encoding *C*. The cortical population activity serves as input to two reinforcement learning modules (right) that control vergence and accommodation commands. For details, see Methods.

Our model is presented with a textured planar object. The object is sampled by the two eyes for ten iterations, constitute one fixation. After each fixation, a new object is presented at a new, random distance. The retinal images are rendered based on the positions of the accommodation, vergence, and object planes (Fig. 3). The inputs are whitened, contrast adjusted by an interocular suppression mechanism, and then binocularly encoded by a population of cortical neurons. The reinforcement learning modules control the retinal input of the next iteration by shifting accommodation and vergence planes along the egocentric axis (Fig. 3).

**Fig. 3.**
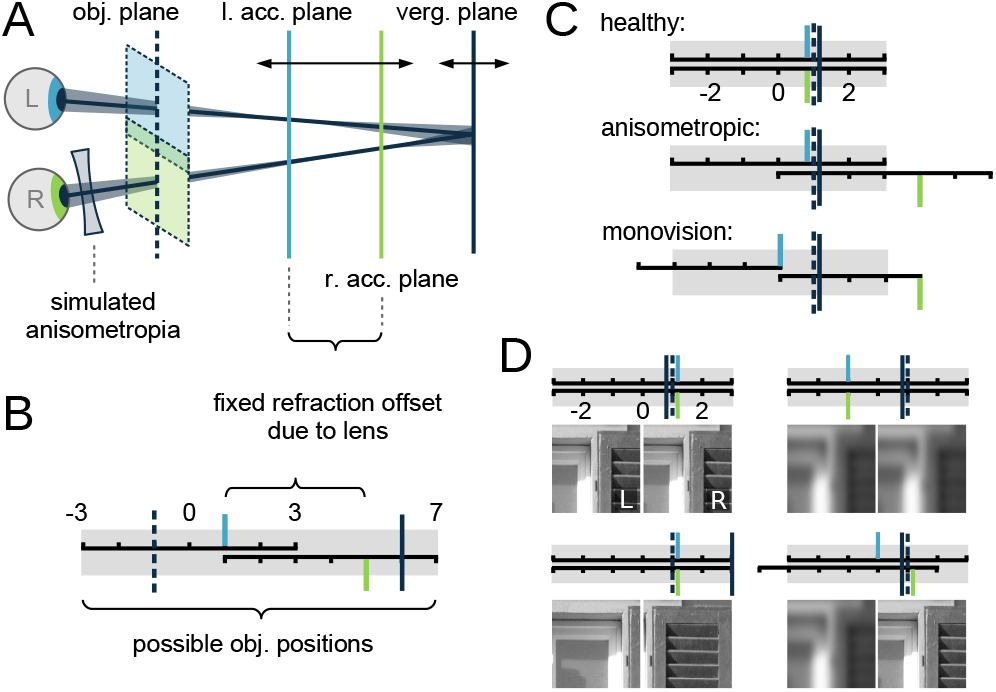
Input sampling from the environment. *(A)* Object position, eye-focus, and eye fixation at different distances are represented as different plane positions. *(B)* Abstraction of *A*. The gray horizontal bar indicates the range where objects are presented during the simulation and also indicates the fixation range, i.e., possible vergence plane positions. Horizontal axes indicate reachable accommodation plane positions for the left (light blue) and right (green) eye. Note that when the stimulus is placed at, e.g., position 0 it cannot be focused by the right eye in this example. Accommodation and vergence errors are measured as the distance between the respective planes and the object position, in a.u. *(C)* Position range of accommodation and vergence planes under different conditions. Same scheme as in *B*. *(D)* Examples of retinal input images for different plane position configurations. For better visibility, disparity shifts and defocus blur are increased compared to actual values.

### Cortical Encoding

In our model, the cortical population activity represents the binocular ‘percept’ based on which behavioral commands are generated (compare Fig. 1, top). The cortical encoding comprises two efficient coders: One for fine details in the foveal region and one for the periphery that receives a low-pass filtered input. Both are implemented using the standard matching pursuit algorithm (26) (see Methods).

To find a set of neurons that best encodes the input, instead of minimizing the conditional entropy directly, we minimize an upper bound, i.e., the average of the encoding error ‖*S*‖^2^ (27):

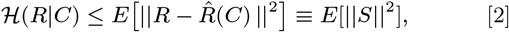

where 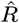 is an estimate of the input *R* based on the activities *c*_*j*_ of cortical neurons with receptive fields *b*_*j*_ (see Methods for details):

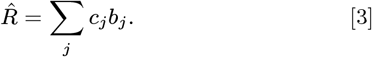

In every iteration, both, activities and receptive fields, adjust online to minimize the encoding error ‖*S*‖^2^ (see Methods). Thus, the receptive fields reflect the stimulus statistics (28) and resemble those of simple cells in the visual cortex (6) (SI Appendix, Fig. S1).

### Vergence Learning

The vergence reinforcement learner also aims to minimize the conditional entropy 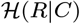, i.e., the encoding error ‖*S*‖^2^. Therefore, vergence movements are favored that produce visual input that can be most accurately encoded with the current set of receptive fields. This leads to a self-reinforcing feedback cycle (Fig. 4*A*). If inputs of a certain disparity can be encoded particularly well, the vergence learner will try to produce visual input that is dominated by this disparity. This will cause even more neurons to become selective for this disparity and make the encoding of this disparity even more efficient (Fig. 4*B*). Thus, an initial bias for, say, small disparities can be magnified until the model always favors input with small disparities and most neurons are tuned to small disparities.

**Fig. 4.**
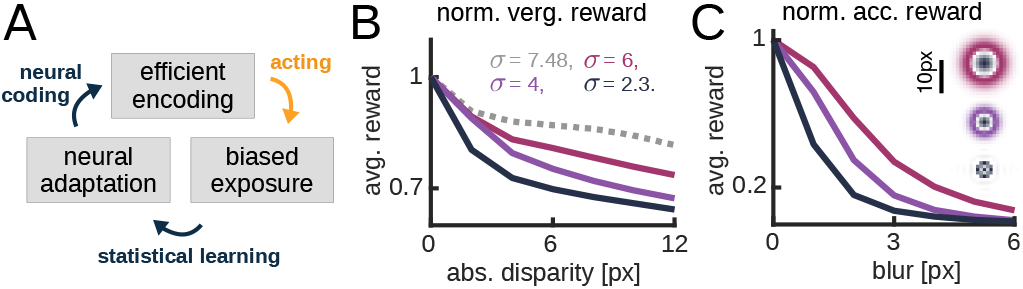
The feedback loop of *Active Efficient Coding* and reward dependencies. *(A)* Positive feedback loop of *Active Efficient Coding*. An efficiently encoded stimulus is preferred over other stimuli (acting). Therefore, the sensory system is more frequently exposed to the stimulus and neural circuits adapt to reflect this overrepresentation (statistical learning) which further increases encoding efficiency (neural coding). *(B)* Normalized vergence reward for different disparity distributions and neural populations. Averaged over 300 textures. The receptive fields of 300 neurons adapted to different distributions of input disparities with color-coded standard deviations. Gray: unbiased/uniform, pink & purple: laplacian distributed, dark blue: model trained under healthy conditions). In each case, stimuli seen at zero disparity produce the highest vergence reward, i.e., the most efficient encoding. This advantage is even more pronounced when small disparities have been encountered more frequently, i.e., for smaller *σ*. *(C)* Normalized accommodation reward for different whitening filters. Zero blur input yields the highest accommodation reward, independent of the size of the whitening filter. However, smaller whitening filters induce a stronger preference for focused input. The smallest filter (dark blue) was used for the simulation (see Methods for details).

### Accommodation Learning

The entropy of the retinal response 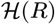 (Eq. 1) is maximized via the accommodation reinforcement learning module. For this, H(R) is approximated by the squared activity of the retinal representation ‖*R*‖^2^ (see Methods). We assume the spatial frequency tuning of retinal ganglion cells to be static and thus independent of the distribution of spatial frequencies in the retinal input as suggested by deprivation experiments (9, 29, 30). However, the exact receptive field shape does not matter for the model to favor focused input (Fig. 4*C*).

### Suppression Model

Interocular suppression is thought to be a central mechanism in amblyopia. We employ a basic interocular suppression model (Fig. 5*A*) to describe dynamic contrast modulation based on the ocular balance of the input encoding. If mostly right (left) monocular receptive fields are recruited during cortical encoding, the contrast of the left(right) eye input becomes suppressed in subsequent iterations. This is in agreement with reciprocal excitation of similarly tuned neurons in visual cortex (31, 32). At the same time, the total input energy is kept balanced to ensure similar activity levels for monocular and binocular visual experience as observed experimentally at high contrast levels (33–35) (see Methods). This leads to a self-reinforcing suppression cycle when left and right eye inputs are dissimilar (Fig. 5*B*).

**Fig. 5.**
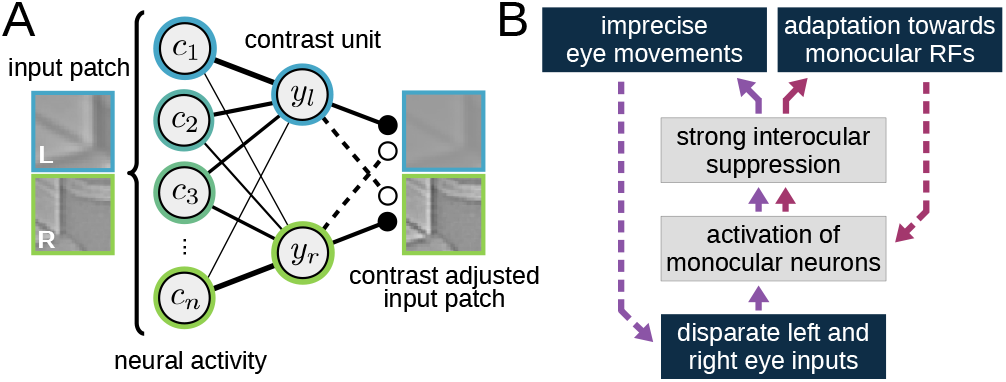
Interocular suppression model. *(A)* When mostly right(left) monocular neurons *c*_*j*_ are activated to encode an input image patch, the right(left) contrast unit *y*_*r*_(*y*_*l*_) is and the left(right) retinal image is suppressed in subsequent iterations. Color hue indicates response selectivity for left eye (blue) or right eye (green). Dashed(solid) lines indicate inhibitory(excitatory) interactions. Connection strength is represented line thickness. We model interocular suppression as being scale specific, i.e., when the high-resolution foveal region of the left eye is suppressed, the low-resolution periphery of the left eye may still provide unattenuated input (see Methods). *(B)* Feed-back cycle of the suppression model. Disparate inputs to both eyes lead to preferential recruitment of monocular neurons, which results in interocular suppression inducing competition between the eyes. This impedes precise vergence eye movements and exacerbates disparate input (purple, left cycle).On a slower timescale, receptive fields (RFs) adapt to suppression by becoming more monocular, which makes future suppression more likely (red, right cycle). Dashed lines indicate feedback that affects future input processing.

## Results

### Active Efficient Coding Leads to Self-calibration of Active Binocular Vision

In the healthy condition without refractive errors, the model learns to perform precise vergence and accommodation eye movements (Fig. 6*A* & SI Appendix, Fig. S2*A*). The object is continuously tracked by the eyes (SI Appendix, Fig. S3*A*) and most neurons develop binocular receptive fields (SI Appendix, Fig. S1). This is not due to artificially introducing a bias for zero disparity during initialization of the model. When receptive fields are adapted to a uniform input disparity distribution, the encoding of zero disparity input is still most efficient (Fig. 4*B*). Due to the overlap of the left and right eye visual field, the information contained in the retinal response 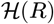 is smallest for zero disparity, when the images projected onto the two eyes maximally overlap. Thus, even an unbiased encoder that can encode inputs of all disparities equally well, will tend to encode zero disparity input more accurately, because such input contains less information. This bootstraps the positive feedback loop of *Active Efficient Coding* (Fig. 4*A*, SI Appendix Fig. S4).

**Fig. 6.**
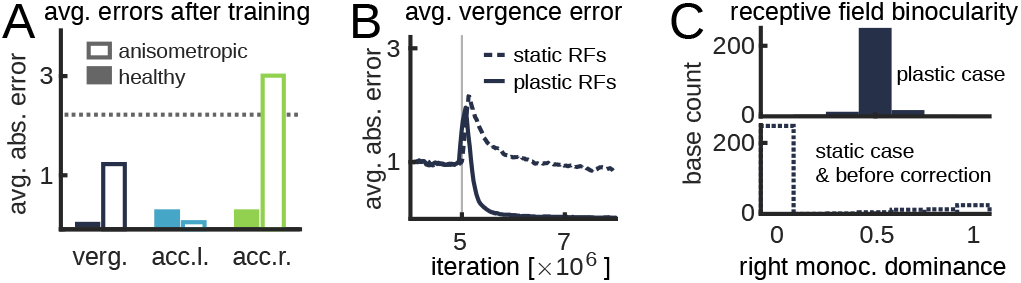
Model performance. *(A)* Average absolute vergence and accommodation errors after training under healthy and anisometropic conditions. The dashed line indicates the expected average vergence error when accommodation planes are moved randomly under healthy conditions. *(B)* Vergence performance of formerly anisometropic model after correction of all refractive errors at iteration 5×10^6^ (vertical gray line). The (dotted)solid line indicates the model with (non-)plastic receptive fields (RFs). The initial increase in the vergence error is due to the recalibration of the reinforcement learning module. *(C)* Histogram of foveal receptive fields binocularity as measured by the right monocular dominance *d*_(·,*r*)_ before and after refractive error correction (see Methods for details).

Accommodation performance becomes highly accurate as well. This is due to the edge-enhancing nature of retinal ganglion cell receptive fields. With their center-surround shape, they are selective for sharp contrasts and respond poorly when out of focus input is presented (Fig. 4*C*). For sharper input, the range of responses across the population, and thus the response entropy, increases (SI Appendix, Fig. S5 & S6).

Furthermore, accurate accommodation is achieved without obvious sign cues: in our simplified visual environment, defocus blur is independent of whether an eye focuses behind or in front of the object. Also, neither chromatic nor other higher-order aberrations are provided in our model, which could help to steer focus in the right direction (36, 37). Instead, the model learns to infer the sign of defocus from disparity cues (SI Appendix, Fig. S7). We further examined this entanglement under abnormal input conditions, e.g. when simulated lenses were placed in front of the eyes of an agent trained under healthy conditions. We find the responses of the model to qualitatively agree with experimental results (38, 39) (SI Appendix, Fig. S8).

### Anisometropia drives model into amblyopic state

To test how the model evolves under abnormal rearing conditions, we simulated an anisometropic case by adding a simulated lens in front of the right eye such that it became hyperopic and was unable to focus objects at close distances (Fig. 3*C*, center). Therefore, unlike the healthy case, where neither eye is favored over the other, in the anisometropic case, the impaired eye receives systematically more defocused input. Cortical receptive fields reflect this imbalance and become more monocular, favoring the unimpaired eye (compare Fig. 6*C*, bottom and Fig. S1, top, center). The combined effect of imbalanced input and adapting receptive fields results in a vicious cycle that drives the model into an amblyopia-like state (Fig. 5*B*). Foveal input from the hyperopic eye becomes actively suppressed (SI Appendix, Fig. S9*A*) while the low-resolution peripheral input is unaffected and still provides binocular information such that a coarse control of vergence is maintained (Fig. 6*A* and SI Appendix, Fig. S2*B* & S3*B*). This results in stable binocular receptive fields in the periphery (Fig. S1, bottom, center), which provide enough information for coarse stereopsis as observed in experiments (40–42). Accommodation adapts such that the stimulus is continuously tracked with the unimpaired eye (SI Appendix, Fig. S3).

When both eyes were similarly impaired but with opposite sign of the refractive error (Fig. 3*C*, bottom), receptive fields still become more monocular but no eye is preferred (SI Appendix, Fig. S1, top, right). As a result, the relatively more myopic eye is used for near and the relatively less myopic eye for distant vision (SI Appendix, Fig. S10*A*) and the respective other, defocused eye is suppressed. At intermediate ranges, the stimulus history determines which eye gets recruited (SI Appendix, Fig. S10*B* & *C*). This configuration is similar to monovision, which results from a treatment method for presbyopia, where the ametropic condition is achieved via optical lenses or surgery (43).

### Early but not late refractive correction rescues binocular vision

To test if the anisometropic model can recover from amblyopia upon correction of the refractive error, we first trained a fully plastic model under anisometropic conditions until it had converged to the amblyopic state. Then, all refractive errors were corrected. When the receptive fields were fixed after the refractive error was corrected, receptive fields remained monocular and the model did not recover from the amblyopic state. Instead, it maintained a high level of vergence error (Fig. 6*B*). In contrast, when receptive fields remained plastic and could adapt to the changed input statistics, the vergence error decreased (Fig. 6*B*) and the strong suppression of the formerly impaired eye was restored to lower values (SI Appendix, Fig. S9*B*). This was due to a shift from monocular to binocular receptive fields as a result of the changed input statistics (Fig. 6*C*). This is in line with a large body of evidence suggesting that limited cortical plasticity in adults prevents recovery from amblyopia after the correction of refractive errors (10, 20, 21). Furthermore, it predicts that therapies reinstating visual cortex plasticity should be effective.

## Discussion

We have shown how simultaneously optimizing both behavior and encoding for efficiency leads to the self-calibration of active binocular vision. Specifically, our model, which is based on a new formulation of the *Active Efficient Coding* theory, accounts for the simultaneous development of vergence and accommodation.

Previous computational models have focused on either the development of disparity tuning or the development of vergence and accommodation control, but have failed to capture their rich interdependence (28, 44–46). For example, a model by Hunt et al. (28) explained how disparity tuning may emerge through sparse coding and how alternate rearing conditions could give rise to systematic differences in receptive field properties, but their model completely neglected vergence and accommodation behavior. Conversely, others have presupposed populations of cells readily providing error signals for vergence and accommodation control without explaining their developmental origin (44, 46). Therefore, previous models have failed to explain how the visual system solves the fundamental ‘chicken and egg’ of disparity tuning and eye movement control: the development of fine disparity detectors requires the ability to accurately focus and align the eyes, which in turn relies on the ability to detect fine disparities. Our *Active Efficient Coding* model solves this problem through the positive feedback loop between disparity tuning, which facilitates the control of eye movements, and improved accommodation and vergence behavior, which enhances the representation of fine disparities. In the end, the tuning properties of sensory neurons reflect the image statistics produced by the system’s own behavior (47). Under healthy conditions, the model develops accurate vergence and accommodation eye movements. For a simulated anisometropia, however, where one eye suffers from a refractive error while the other eye is unaffected, it develops into an amblyopia-like state with monocular receptive fields and loss of fine stereopsis. Recovery from this amblyopia-like state is only possible if receptive fields in the model remain plastic, matching findings of a critical period for binocular development (10).

An important mechanism in amblyopia is interocular suppression. The simple logic behind the model’s suppression mechanism is that every neuron suppresses input that is incongruent to its own receptive field (34, 35). This implementation proved sufficient to account for the development of an amblyopia-like state, with mostly monocular receptive fields in the representation of the fovea. More sophisticated suppression models could be incorporated in the future (48, 49), but we do not expect them to change the conclusions from the present model. Future work should focus on understanding the principles of interocular suppression within the *Active Efficient Coding* framework. A topic of current interest is how suppression develops during disease and treatment, e.g., with the standard patching method (50). A better understanding of the role of suppression in amblyopia could lead to improved therapies in the future.

While we have focused on the development of active binocular vision including accommodation and vergence control, our formulation of *Active Efficient Coding* is very general and could be applied to many active perception systems across species and sensory modalities. *Active Efficient Coding* is rooted in classic efficient coding ideas (2–6), of which predictive coding theories are special examples (51–53). Classic efficient coding does not consider optimizing behavior, however. Friston’s Active Inference approach does consider the generation of behavior in a very general fashion. There, motor commands are generated to fulfill sensory predictions. In our new formulation of *Active Efficient Coding*, motor commands are learned to maximize the mutual information between the sensory input and its cortical representation. This implies maximizing the amount of sensory information sampled from the environment and avoids the problem of deliberately using accommodation to defocus the eyes, or closing the eyes altogether, to make the sensory input easy to encode and/or predict.

## Materials and Methods

### Input Image Rendering

We used 300 grayscale converted natural images of the ‘man-made’ category from the McGill Database (54). One image was presented at a random position (see Fig. s3*A* and *B*) during one fixation, i.e., 10 subsequent iterations, before the next image and position were randomly selected for the next fixation.

For every distance unit between vergence and object plane, the left(right) eye image was shifted 1px to the left(right). This resulted in a disparity of 2 px per distance unit. A Gaussian blur filter was applied to the left and the right eye image where the standard deviations depended linearly on the distance between object and accommodation planes. 1 a.u. distance equals 0.8 px of standard deviation (SI Appendix text, Fig. S11 & Fig. S12). Errors were measured in a.u. as distances to the object plane. For the foveal(peripheral) scale, two retinal images of size 72 × 72(160 × 160) pixels were cropped from the center of the original image (see SI Appendix, Fig. S13).

### Input processing

The left and right retinal input images were whitened as described by Olshausen and Field (6) (see SI Appendix for details). For each scale, images were cut and merged into 81 binocular patches of size 2×8×8 px, where the peripheral scale was downsampled with a Gaussian pyramid by a factor of 4 (see SI Appendix, Fig. S13). The whitened retinal patches *R*\* were normalized to zero mean intensity and subsequently contrast adjusted via the interocular suppression mechanism (see below). The contrast adjusted patches *R* were encoded with the matching pursuit algorithm (26).

For each patch, we recruited *N* = 10 out of 300 cortical neurons to most efficiently encode the image. The cortical response *C* = (*c*_1_, ⋯, *c*_300_) was determined via an iterative process, where the activities of neurons that were not selected for encoding remained zero. In the first encoding step, *n* = 1, the neuron whose receptive field *b*_*j*_ was most similar to the retinal input was selected.

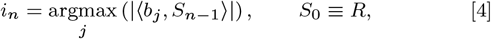

where the similarity between a receptive field *b*_*j*_ and retinal input *R* was meassured with the scalar product ⟨*b*_*j*_, *R*⟩. When selecting the next neuron, all information that was already encoded by the first neuron is substraced from the original input *R*:

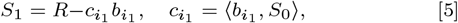

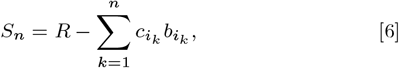

where *S*_1_ is the residual image after the first encoding step and *S*_*n*_ is the generalized residual after the *n*-th encoding step. Subsequent neurons are selected based on the similarity of their receptive fields with the residual according to Eq. 4. By greedily selecting the neuron with maximum response 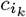 in each encoding step, the reconstruction error ‖*S*_*N*_‖^2^ is minimized, i.e., coding efficiency is maximized.

After encoding, all receptive fields were updated through gradient descent on ‖*S*_*N*_‖^2^ and normalized to unit length. Thus, their tuning reflects the input statistics (SI Appendix, Fig. S1).

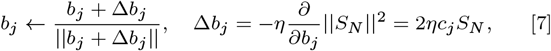

where 2*η* = 5 × 10^−5^ is a learning rate. Each patch of the foveal scale was encoded by a subset of the same 300 neural receptive fields. For the peripheral scale a separate set of 300 neurons was used for encoding (see SI Appendix, Fig. S13). At the beginning of each simulation, all receptive field weights were drawn randomly from a zero-mean Gaussian distribution and subsequently normalized to unit norm.

### Reinforcement Learning

We used two separate natural actor critic reinforcement learners (55, 56) with identical architectures to control the accommodation planes and the vergence plane, respectively. Possible actions *a* correspond to shifts in the respective plane positions: *a* ∈ {−2, −1, 0, 1, 2} (compare Fig. 3). The state information vector comprises the patch-averaged squared responses of the cortical neurons:

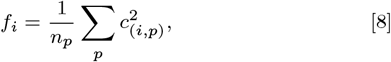

where *c*_(*i,p*)_ is the activity of neuron *i* after encoding patch *p* and *n*_*p*_ = 81 is the number of patches per scale. Therefore, *f*_*i*_ is spatially invariant, due to averaging over patches, and does not depend on the polarity of the input, due to the squaring. This is similar to the properties of complex cells in primary visual cortex (57, 58). After they were normalized to unit norm, the peripheral and the foveal scale state vector are concatenated into the combined state vector *f* of size 300 × 2. The next action *a* is choosen with probability *π*_*a*_(*f*) and the state value is estimated as *V*(*f*):

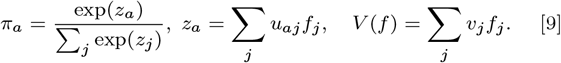

The weights *u*_*aj*_ and *v*_*j*_ are updated via approximate natural gradient descent on an approximation of the temporal difference error *δ* (algorithm 3 in (56)):

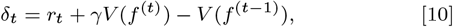

where *r*_*t*_ is the accommodation or vergence reward and *γ* = 0.6 is the temporal discounting factor. At the beginning of each simulation, all network weights were initialized randomly.

### Approximating Mutual Information

Rewards for the reinforcement learners are based on the squared response after whitening and cortical encoding, respectively. Together this can be understood as an empirical estimate of the mutual information between the whitened response *R* and cortical response *C*:

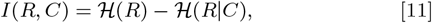

where the conditional entropy 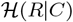, is upper bounded by the reconstruction error ‖*S*‖^2^ (see Eq. 2 & SI Appendix). Due to the ‘energy conservation’ property of the matching pursuit algorithm (26), the energy of the residual image is equal to the energy of the retinal representation minus the energy of the cortical representation (see SI Appendix), i.e.,

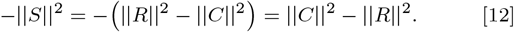

Therefore, we take the difference between cortical and retinal response energy as the reward for the vergence learner.

For the accommodation learner, we maximize the entropy of the whitened retinal response 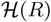. We take each entry of *R* as an independent sample of the same underlying random variable and estimate the entropy of its probability distribution. The distribution is well approximated by a Laplace distribution, independent of the level of blur in the input (Fig. S5). Therefore, we approximate 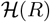 with the entropy of a Laplace distribution with the same standard deviation *σ*_*R*_:

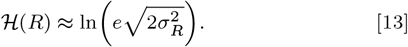

Since the expected squared activity of the retinal representation *E*(‖*R*‖^2^) is equal to the variance 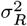, it is also a monotonic function of the entropy 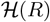. More generally, since the retinal response has bounded support and its probability distribution is unimodal and Lipschitz-continuous, the variance 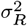 is a monotonic function of a lower bound of the entropy(59). Therefore, we use ‖*R*‖^2^, as an empirical estimate of *E*(‖*R*‖^2^), for the reward of the accommodation reinforcement learning module.

As one would expect for the entropy 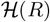, also ‖*R*‖^2^ decreases for increasing input blur. Under the assumption of a flat frequency spectrum after whitening, one finds (see SI Appendix):

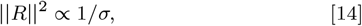

where *σ* is the standard deviation of the Gaussian blur filter that is applied before whitening to simulate defocus blur.

### Reward Normalization

Before being passed to the reinforcement learning agents, the accommodation and the vergence rewards were normalized online to zero mean and unit variance, i.e.,

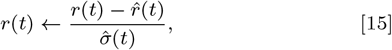

where 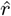 is the exponentially weighted running average of the reward *r* and 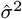 is an online estimate of its variance:

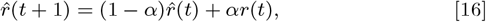

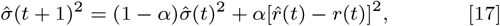

where *α* = 0.001 is an update rate that sets the decay of the exponential weighting (60).

### Suppression Mechanism

There are two separate suppression modules, one per scale, that adjust the contrast of left and right input image (see Fig. 5). We introduce a contrast measure *x*_*k*_, *k* ∈ {*l, r*} that gives an estimate of the amount of left(right) monocular input over the previous iterations.

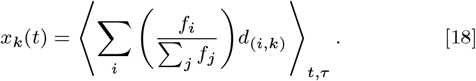

The monocular dominance of each neuron *d*_(*i,k*)_ is weighted with its relative patch-averaged squared activation *f*_*i*_/∑_*j*_ *f*_*j*_. Here, ⟨·⟩_*t,τ*_ is the exponential moving average over time with decay constant *τ* = 10 (see SI Appendix). The monocular dominance is defined as:

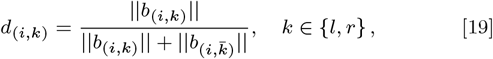

where *b*_(*i,k*)_ is the left/right monocular subfield of neuron *i*. The contrast estimate *x*_*k*_ of the left and right subfield of the input image is separately processed by two contrast units

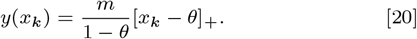

As the contrast estimate *x*_*k*_ crosses the threshold *θ*, the output *y*(*x*_*k*_) increases from zero until it saturates at *m*. We chose the threshold *θ* = 0.6, just above perfect binocular input at *x*_*k*_ = 0.5 to provide some margin before the self-reinforcing feedback loop becomes active (Fig. 5B). Further, we set the saturation *m* = 0.8 to prevent total suppression of one eye. Finally, the subsequent input sub-patches for the cortical coder are adjusted to

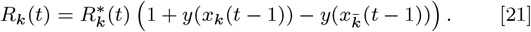

Note that *I*(*R, C*) = *I*(*R*\*, *C*), since *R* is homeomorphic to *R*\* (61). Therefore, in our theoretical framework, we do not distinguish between the contrast adjusted and the raw retinal response. For the model implementation, the contrast adjusted retinal response *R* is employed. See SI Appendix for additional detail.

### Software and Documentation

Documented MATLAB code of the model is available in ModelDB under accession number 261483.

## ACKNOWLEDGMENTS

This work was supported by the German Federal Ministry of Education and Research under Grants 01GQ1414 and 01EW1603A, the Hong Kong Research Grants Council under Grant 618713, and the Quandt Foundation. We thank Maria Fronius and Rowan Candy for numerous discussions.

## Notes

The authors declare no conflict of interest.

http://modeldb.yale.edu/261483

## References

1. Clark DD, Sokoloff L (1999) Circulation and energy metabolism of the brain in Basic Neurochemistry: Molecular, Cellular and Medical Aspects, eds. Siegel GJ, Agranoff BW, Albers RW, Fisher SK, Uhler MD. (Lippincott-Raven, Philadelphia), p. 637–670.

2. Barlow HB,, et al. (1961) Possible principles underlying the transformation of sensory messages. Sensory communication 1:217–234.

3. Simoncelli EP, Olshausen BA (2001) Natural image statistics and neural representation. Annual review of neuroscience 24(1):1193–1216.

4. Lewicki MS (2002) Efficient coding of natural sounds. Nature neuroscience 5(4):356.

5. Atick JJ, Redlich AN (1992) What does the retina know about natural scenes? Neural computation 4(2):196–210.

6. Olshausen BA, Field DJ (1997) Sparse coding with an overcomplete basis set: A strategy employed by v1? Vision research 37(23):3311–3325.

7. Wiesel TN, Hubel DH (1963) Single-cell responses in striate cortex of kittens deprived of vision in one eye. Journal of neurophysiology 26(6):1003–1017.

8. Hirsch HV, Spinelli D (1970) Visual experience modifies distribution of horizontally and vertically oriented receptive fields in cats. Science 168(3933):869–871.

9. Movshon JA, Van Sluyters RC (1981) Visual neural development. Annual review of psychology 32(1):477–522.

10. Espinosa JS, Stryker MP (2012) Development and Plasticity of the Primary Visual Cortex. Neuron 75(2):230–249.

11. Zhao Y, Rothkopf CA, Triesch J, Shi BE (2012) A unified model of the joint development of disparity selectivity and vergence control in Development and Learning and Epigenetic Robotics (ICDL), 2012 IEEE International Conference on. (IEEE), pp. 1–6.

12. Teulière C, et al. (2015) Self-calibrating smooth pursuit through active efficient coding. Robotics and Autonomous Systems 71:3–12.

13. Gibson JJ (1966) The Senses Considered as Perceptual Systems. (Boston, USA: Houghton Mifflin).

14. Friston K, Thornton C, Clark A (2012) Free-energy minimization and the dark-room problem. Frontiers in psychology 3:130.

15. Sims A (2017) The problems with prediction in Philosophy and Predictive Processing, eds. Metzinger TK, Wiese W. (MIND Group, Frankfurt am Main).

16. Tondel GM, Candy TR (2008) Accommodation and vergence latencies in human infants. Vision research 48(4):564–576.

17. Blakemore C, Van Sluyters RC (1974) Experimental analysis of amblyopia and strabismus. The British journal of ophthalmology 58(3):176.

18. Kenyon RV, Ciuffreda K, Stark L (1980) Dynamic vergence eye movements in strabismus and amblyopia: symmetric vergence. Investigative ophthalmology & visual science 19(1):60–74.

19. Manh V, Chen AM, Tarczy-Hornoch K, Cotter SA, Candy TR (2015) Accommodative performance of children with unilateral amblyopia. Investigative ophthalmology & visual science 56(2):1193–1207.

20. Levi DM, Knill DC, Bavelier D (2015) Stereopsis and amblyopia: a mini-review. Vision research 114:17–30.

21. Stryker MP, Loewel S (2018) Amblyopia: New molecular/pharmacological and environmental approaches. Visual neuroscience 35.

22. Cotter SA, Group PEDI,, et al. (2006) Treatment of anisometropic amblyopia in children with refractive correction. Ophthalmology 113(6):895–903.

23. Steele AL, et al. (2006) Successful treatment of anisometropic amblyopia with spectacles alone. Journal of American Association for Pediatric Ophthalmology and Strabismus 10(1):37–43.

24. Zhu Q, Triesch J, Shi BE (2017) Joint learning of binocularly driven saccades and vergence by active efficient coding. Frontiers in neurorobotics 11:58.

25. Klimmasch L, Lelais A, Lichtenstein A, Shi BE, Triesch J (2017) Learning of active binocular vision in a biomechanical model of the oculomotor system in 2017 Joint IEEE International Conference on Development and Learning and Epigenetic Robotics (ICDL-EpiRob). (IEEE), pp. 21–26.

26. Mallat SG, Zhang Z (1993) Matching pursuits with time-frequency dictionaries. IEEE Transactions on signal processing 41(12):3397–3415.

27. Cover TM, Thomas JA (2006) Elements of information theory. (John Wiley & Sons).

28. Hunt JJ, Dayan P, Goodhill GJ (2013) Sparse Coding Can Predict Primary Visual Cortex Receptive Field Changes Induced by Abnormal Visual Input. PLoS Computational Biology 9(5).

29. Sherman SM, Stone J (1973) Physiological normality of the retina in visually deprived cats. Brain Research 60(1):224–230.

30. Movshon JA, et al. (1987) Effects of early unilateral blur on the macaque’s visual system. iii. physiological observations. Journal of Neuroscience 7(5):1340–1351.

31. Yoshioka T, Blasdel GG, Levitt JB, Lund JS (1996) Relation between patterns of intrinsic lateral connectivity, ocular dominance, and cytochrome oxidase-reactive regions in macaque monkey striate cortex. Cerebral Cortex 6(2):297–310.

32. Iacaruso MF, Gasler IT, Hofer SB (2017) Synaptic organization of visual space in primary visual cortex. Nature 547(7664):449–452.

33. Moradi F, Heeger DJ (2009) Inter-ocular contrast normalization in human visual cortex. Journal of Vision 9(3):13–13.

34. Said CP, Heeger DJ (2013) A model of binocular rivalry and cross-orientation suppression. PLoS computational biology 9(3):e1002991.

35. Carandini M, Heeger DJ (2012) Normalization as a canonical neural computation. Nature Reviews Neuroscience 13(1):51.

36. Chen L, Kruger PB, Hofer H, Singer B, Williams DR (2006) Accommodation with higher-order monochromatic aberrations corrected with adaptive optics. Journal of the Optical Society of America. A, Optics, image science, and vision 23(1):1–8.

37. Zannoli M, Love GD, Narain R, Banks MS (2016) Blur and the perception of depth at occlusions. Journal of Vision 16(6):17–17.

38. Bharadwaj SR, Candy TR (2009) Accommodative and vergence responses to conflicting blur and disparity stimuli during development. Journal of vision 9(11):4.1–18.

39. Bharadwaj SR, Candy TR (2011) The effect of lens-induced anisometropia on accommodation and vergence during human visual development. Investigative Ophthalmology and Visual Science.

40. Holopigian K, Blake R, Greenwald MJ (1986) Selective losses in binocular vision in anisometropic amblyopes. Vision research 26(4):621–630.

41. Babu RJ, Clavagnier S, Bobier WR, Thompson B, Hess RF (2017) Regional extent of peripheral suppression in amblyopia. Investigative Ophthalmology and Visual Science 58(4):2329–2340.

42. Bradley A, Freeman RD (1981) Contrast sensitivity in anisometropic amblyopia. Investigative Ophthalmology and Visual Science 21(3):467–476.

43. Evans BJ (2007) Monovision: A review. Ophthalmic and Physiological Optics 27(5):417–439.

44. Schor CM, Alexander J, Cormack L, Stevenson S (1992) Negative feedback control model of proximal convergence and accommodation. Ophthalmic and Physiological Optics 12(3):307–318.

45. Hoyer PO, Hyvärinen A (2000) Independent component analysis applied to feature extraction from colour and stereo images. Network: computation in neural systems 11(3):191–210.

46. Gibaldi A, Chessa M, Canessa A, Sabatini SP, Solari F (2010) A cortical model for binocular vergence control without explicit calculation of disparity. Neurocomputing 73(7-9):1065–1073.

47. Gibaldi A, Canessa A, Sabatini SP (2017) The active side of stereopsis: Fixation strategy and adaptation to natural environments. Scientific reports 7:44800.

48. Hess RF, Thompson B (2015) Amblyopia and the binocular approach to its therapy. Vision Research 114:4–16.

49. Hallum LE, et al. (2017) Altered balance of receptive field excitation and suppression in visual cortex of amblyopic macaque monkeys. The Journal of Neuroscience 37(34):8216–8226.

50. Kehrein S, Kohnen T, Fronius M (2016) Dynamics of interocular suppression in amblyopic children during electronically monitored occlusion therapy: first insight. Strabismus 24(2):51–62.

51. Rao RP, Ballard DH (1999) Predictive coding in the visual cortex: a functional interpretation of some extra-classical receptive-field effects. Nature neuroscience 2(1):79.

52. Friston K (2010) The free-energy principle: a unified brain theory? Nature reviews neuroscience 11(2):127.

53. Palmer SE, Marre O, Berry MJ, Bialek W (2015) Predictive information in a sensory population. Proceedings of the National Academy of Sciences 112(22):6908–6913.

54. Olmos A, Kingdom FA (2004) A biologically inspired algorithm for the recovery of shading and reflectance images. Perception 33(12):1463–1473.

55. Sutton RS, Barto AG (2018) Reinforcement learning: An introduction. (MIT press).

56. Bhatnagar S, Sutton R, Ghavamzadeh M, Lee M (2009) Natural actor-critic algorithms. Automatica 45(11).

57. Hubel DH, Wiesel TN (1962) Receptive fields, binocular interaction and functional architecture in the cat’s visual cortex. The Journal of physiology 160(1):106–154.

58. Adelson EH, Bergen JR (1991) The Plenoptic Function and the Elements of Early Vision, ed. M. Landy and J. A. Movshon. (The MIT Press), pp. 3–20.

59. Chung HW, Sadler BM, Hero AO (2017) Bounds on variance for unimodal distributions. IEEE Transactions on Information Theory 63(11):6936–6949.

60. Finch T (2009) Incremental calculation of weighted mean and variance. University of Cambridge Computing Service.

61. Kraskov A, Stögbauer H, Grassberger P (2004) Estimating mutual information. Phys. Rev. E 69(6):066138.

