## Supplementary Information for "Active Efficient Coding Explains the Development of Binocular Vision and its Failure in Amblyopia"

#### This PDF file includes:

Supplementary text
Figs. S1 to S13
References for SI reference citations

### Supporting Information Text

**Input Rendering.** For every unit of distance between the vergence and the object plane, a retinal disparity of 2 px was introduced. This results in a 2 px shift between the left and right eye input. For example, if the object plane lies in front of the vergence plane, the left eye's field of view was shifted one pixel to the left and the right eye's field of view was shifted one pixel to the right (see Main text, Fig. 3). Here, we implicitly assumed that the brain operates in units of disparity on the retina and not in units of physical distance between vergence and object plane. This justifies our crude approximation of a linear relationship between plane distance and disparity that only holds for small vergence errors.

To model defocus blur, a gaussian blur filter was applied to the respective eye's input. Its standard deviation was set to 0.8 px times the units of distance between the accommodation plane and the object plane (compare Fig. S11*A*). Accordingly, the filter diameter per unit of distance is approximately 2 px.

In the real visual system the relationship between disparity and defocus blur is not arbitrary but depends on the interocular distance and the pupil size. The average pupile size in humans is approximately $p = 0.3$ cm (1). The average interocular distance (IOD) is roughly 6 cm (2). Then, the ratio between disparity and defocus blur per distance unit of focus or fixation error (compare Main text Fig. 3) is IOD$/p = 20$ (see Fig. S12). This is larger than in our model, were the ratio is approximately 1, with 2 px of disparity shift corresponding to defocus blur that averages over approximately 2 px (compare Fig. S12: $s_1 - s_2$ corresponds to one distance unit of error, $\delta_d' = 2$ px disparity shift and $\delta_b \approx 2$ px defocus blur).

Originally, we chose the ratio such that the differential repercussions of anisometropia on high and low-resolution visual processing becomes apparent. In our model, an anisometropic case elicits suppression in the fovea while the lower-resolution periphery is unaffected (Fig. S9), as multiple experimental studies suggest (3–6). This requires that the defocus blur is large enough to affect the high-resolution foveal scale. Due to rasterization in our input rendering model, the standard deviation of defocus blur cannot be much smaller than one pixel per unit error before the gaussian blur filter becomes too localized to have any noticeable blurring effect. At the same time, defocus blur must be small enough to not affect input processing in the low-resolution peripheral scale. This depends critically on the choice of the resolution for the different scales. The size of receptive fields in the visual cortex increases continuously with eccentricity; by factors $> 100$ between the fovea and the far periphery (7–9). Therefore, our choice for the resolutions of the two different scales (compare Fig. S13) is to some degree arbitrary and the ratio between disparity and defocus is not meaningfully constrained, since the input textures are uncalibrated. For example, we could reinterpret the 2 px of disparity shift per unit error as a 20 px shift of a 20 times larger input texture, but processed on a 20 times lower-resolution scale. In this interpretation, the required ratio is matched to the biological values. We demonstrated in previous work that earlier *Active Efficient Coding* models can operate in more realistic simulated environments (10, 11) and even on real-world robotic platforms (12, 13). In summary, the chosen parameters allow us to demonstrate a qualitative difference between foveal and peripheral processing but do not allow a quantitative comparison to biological data.

**Whitening.** The receptive field shape of retinal ganglion cells has been interpreted as the result of maximizing coding efficiency (14, 15). With their center-surround shape, they reduce the redundancy of natural images (see (16), p.137 for a detailed tutorial). The optimal receptive field transforms the input such that the amplitude spectrum of the Fourier transformation becomes flat (Fig. S6), similar to the frequency spectrum of white noise. Therefore, the receptive field resembles a whitening filter (14).

As in (15), we model the retinal ganglion cell population activity as the image pixel intensities after convolving the sensory input $I$ with a whitening filter $W$. The convolution is performed as a multiplication in Fourier space:

$$R^* = \mathcal{F}^{-1}[\mathcal{F}(W)\mathcal{F}(I)],$$

where $(\mathcal{F}^{-1})\mathcal{F}$ is the (inverse) Fourier transform and $R^*$ is the whitened retinal response before contrast adjustment. We define the whitening filter $W$ in Fourier space as (15):

$$\mathcal{F}(W)(\omega_x, \omega_y) = \rho^a \exp[-(\rho/\rho_0)^\kappa], \quad \rho = \sqrt{\omega_x^2 + \omega_y^2},$$

where $\rho$ is the absolute frequency and $a$ is the decay exponent of the input spectrum (compare Fig. S6). $\rho_0 = 0.4$ c/px (cycles per pixel) defines the cut-off frequency to prevent amplification of high frequency noise (14, 15), while $\kappa = 4$ sets the decay speed for frequencies larger than the cut-off. These are the same parameters as in (15), except that we set $a = 1.27$ to achieve a flat frequency spectrum for our image data set (Fig. S6). However, our model is robust with respect to the exact choice of the whitening filter (compare Main text, Fig. 4*C*).

**Suppression Mechanism.** The suppression mechanism introduces competition between the two eyes mediated by neurons with predominantly monocular receptive fields. In essence, we postulate that each neuron tries to maximize its firing by enhancing input for which it is selective while suppressing competing activity. This is motivated by the connectivity of pyramidal cells in the visual cortex. These cells preferentially excite other pyramidal cells with similar tuning properties (17, 18). Combined with a normalization operation (19, 20), the net effect is the suppression of incongruent information and enhancement of congruent information. This is also a common mechanism in models of binocular rivalry (21, 22).

In our model, we apply this principle by enhancing input from one or the other eye depending on the activities and receptive field binocularities of all cortical neurons. If one neuron with a left dominant receptive field is active, in the next iteration, the contrast of the left image is increased and the contrast in the right image is decreased. This bears a suppressive impact on neurons tuned to a different eye while responses of neurons tuned to the same eye are enhanced.

In a first step, we quantify the binocularity of each neuron $i$ via the monocular dominances $d_{(i,k)}$ of the receptive subfield $b_{(i,k)}$ of the left $(k = l)$ and right $(k = r)$ eye:

$$d_{(i,k)} = \frac{||b_{(i,k)}||}{||b_{(i,k)}|| + ||b_{(i,\bar{k})}||}, \quad k \in \{l, r\}\,. \tag{1}$$

Note that dominance levels lie between 0 and 1, where the two extremes describe monocular neurons tuned to opposite eyes. For $d_{(i,k)} = 0.5$ the neuron is assumed to be perfectly binocular and should suppress neither of the two eyes. In a simplified scenario, the patch averaged activity of a single neuron $\langle c_i^2 \rangle_p$ affects the left eye's whitened retinal response $R_l^*$ based on its monocular dominance $d_{(i,k)}$ as follows:

$$R_l = R_l^* \left(1 + \langle c_i^2 \rangle_p [d_{(i,l)} - d_{(i,r)}]\right),$$

where $R_l$ is the contrast adjusted retinal response of the left eye. $\langle c_i^2 \rangle_p$ can be understood as a model for the response of a complex cell in visual cortex (23, 24), as its activity is spatially invariant, due to averaging over patches (see Methods section of the main manuscript). For better readability, we omit the notation for averaging over all patches in the following. We model suppression and enhancement as multiplicative effects that effectively decrease or increase the contrast of the left or right whitened retinal responses $R_l^*$ and $R_r^*$. Here, $(d_{(1,l)} - d_{(1,r)})$ can be thought of as a binocularity index between $-1$ for right monocular receptive fields and 1 for left monocular receptive fields. For balanced, binocular receptive fields we get $d_{(i,l)} - d_{(i,r)} = 0$ and, therefore, no contrast adjustment of the input. The magnitude of suppression or enhancement is determined by the activation of each neuron $c_i$. In the case of multiple neurons, we take a normalized weighted average:

$$R_l^* \left(1 + \sum_i \left(\frac{c_i^2}{\sum_j c_j^2}\right) d_{(i,l)} - \sum_i \left(\frac{c_i^2}{\sum_j c_j^2}\right) d_{(i,r)}\right) \equiv R_l^*(1 + x_l - x_r), \tag{2}$$

where we split up the suppression and the enhancement term and defined the left and right contrast measure $x_l$ and $x_r$. Due to the normalization via $1/\sum_j c_j^2$, $(x_l - x_r)$ is the activity weighted average binocularity with values between $-1$ and 1. This can be interpreted as a contrast measure that is not based on the input representation but the cortical encoding (compare Fig. S9). As such, it represents the percept of the agent based on the activation of its receptive fields. For example, when there are almost exclusively left monocular receptive fields, i.e., $d(i, r) \approx 0$ for all neurons $i$, the contrast for the right eye is perceived to be very low, independent of the actual input $R_r^*$. As a result, the right eye's input $R_r^*$ gets suppressed and is not encoded at the cortical stage. Due to this suppression, receptive fields adapt towards a left-dominated sensory input and become left-dominated themselves. This leads to even stronger suppression (compare suppressive feedback loop in Main text, Fig. 5*B*).

In the case of exclusively binocular bases, i.e., $d(i, k) = 0.5\ \forall\ i, k$, this feedback loop can be initiated for small contrast imbalances between the left and the right eye: When slightly left monocular input is encoded, the right eye becomes slightly suppressed which results in slightly more left monocular receptive fields and, therefore, slightly more right eye suppression in the future (Main text, Fig. 5*B*). This means the system is unstable and can not maintain binocular vision unless the input is perfectly balanced. To make it more robust, we extended the model as follows.

First, we set the suppression for small imbalances to zero. We recast Eq. 2 as:

$$R_l^*(1 + x_l - x_r) = \begin{cases} R_l^*(1 + 2(x_l - 0.5)), & \text{if} \quad x_l > 0.5 \\ R_l^*(1 - 2(x_r - 0.5)), & \text{if} \quad x_r < 0.5 \end{cases} = R_l^*\left(1 + 2[x_l - 0.5]_+ - 2[x_r - 0.5]_+\right),$$

where we used $x_r = 1 - x_l$ and therefore $x_l > 0.5 \Leftrightarrow x_r < 0.5$. Now each term can be interpreted as the output of a rectified linear "contrast unit" $y$ with throshold $\theta = 0.5$, saturation $m = 1$ and non-linearity $[\cdot]_+ \equiv max(0, \cdot)$. Then, the final suppression mechanism can be expressed as:

$$R_l(t) = R_l^*(t)\left(1 + y(x_l(t-1)) - y(x_r(t-1))\right), \qquad y(x) = \frac{m}{1-\theta}[x - \theta]_+.$$

Suppression starts as soon as $x_l > \theta$. We allow for some deviation from perfect balance at $x = 0.5$ by choosing the threshold as $\theta = 0.6$. This leaves some margin for imbalance between left and right eye input before suppression sets in. The saturation $m$ defines the maximum suppression. When all activity is concentrated at neurons with maximal left dominant receptive fields, i.e., $x_l = 1$, then the contrast of the right eye can be suppressed to a fraction $1 - y(x_l) = 1 - m$ of its original level. If, due to some impairment, the right eye is at a disadvantage and the maximal suppression $m = 1$, then, due to the suppressive feedback loop (Main text, Fig. 5*B*), the right eye input can become completely suppressed and receptive fields adjust such that the right subfield approaches zero. Then, even when the initial impairment is removed, visual input is encoded with exclusively left monocular neurons (compare Eq. 1) and input from the right eye is ignored. In contrast, many neurons in the monocular regions of the visual cortex in cats remain responsive after monocular deprivation (25). In macaques, even in the most severe cases of monocular deprivation, one finds neurons that are sensitive for binocular input (26). Therefore, we limit the maximal suppression level by setting the saturation of the contrast units to $m = 0.8$.

To further increase stability, we assumed that the contrast adjustments due to suppression last longer than it takes to process one stimulus. This means, if a suppressive stimulus is removed, it takes a few iterations for the suppression to be lifted. Therefore, instead of a contrast estimate based on the current input, we take the average $\bar{x}_k$ over multiple previous contrast measures. This reduces the impact of contrast imbalances, as they get smoothed out over time and are less likely to cross the suppression threshold $\theta$. We make use of the exponential moving average $\langle\cdot\rangle_{t,\tau}$:

$$\bar{x}_k(t) \equiv \langle x_k \rangle_{t,\tau}(t) = \left(1 - \frac{1}{\tau}\right)^t x_k(0) + \frac{1}{\tau}\sum_{j=1}^{t}\left(\left(1 - \frac{1}{\tau}\right)^{t-j} x_k(j)\right).$$

This means each previous contrast meassure $x_k(j)$ is weighted with a factor $(1-1/\tau)^{t-j}$ that decays exponentially for measures further in the past, i.e., for smaller $j$, since $(1-1/\tau) < 1$ for $\tau > 1$. Then, $\bar{x}_k(t)$ can be computed recursively, i.e., by a weighted average of the most recent measure $x_k(t)$ and the moving average of the previous timestep $\bar{x}_k(t-1)$:

$$\bar{x}_k(t) = \left(1 - \frac{1}{\tau}\right)\bar{x}_k(t-1) + \frac{1}{\tau}x_k(t).$$

Here, $\tau$ sets the timescale how fast the weighting of previous measures decreases. We choose the timescale to be on the order of one fixation, i.e., $\tau = 10$ iterations in our model. However, this does not mean that the amblyopia-like state our model develops in the anisometropic case can be lifted rapidly, i.e., on the timescale of one fixation. We find that the exact choice of $\tau$ does not matter for the development of the amblyopia-like state. In the anisometropic case, the suppression is due to a lack of neurons that are tuned to the suppressed eye (compare Fig. S1, center top, and Main text, Fig. 6*C*, bottom). This leads to strong suppression, even when balanced input is provided. The receptive fields are adjusted on a much slower timescale, such that only prolonged binocular input can lift suppression. In contrast, in the monovision case, there are mostly monocular neurons tuned to one or the other eye (Fig. S1, top, right), such that suppression can switch between the eyes within just one fixation (Fig. S10).

In our model, there are two separate suppression mechanisms, one for the foveal and one the peripheral scale. This enables the foveal scale to become suppressed while the peripheral scale remains binocular. For example, in anisometropia, high spatial frequencies may be attenuated in one eye. Therefore, on the foveal scale, left and right eye input are vastly different and suppression sets in (compare Main text, Fig. 5*B*), while the lack of high frequencies may be unrecognizable at the lower-resolution peripheral scale, where higher frequencies are not detected (compare Fig. S9*A*). This is in accordance with experimental evidence that reports functional stereopsis for lower spatial frequencies or in the peripheral visual field of patients with anisometropic amblyopia (3, 4, 6).

**Energy Conservation Property of the Matching Pursuit Algorithm.** The $n$-th encoding step of the matching pursuit algorithm (27) can be expressed as (compare Methods section):

$$S_n = R - \sum_{k=1}^{n} c_{i_k} b_{i_k}, \quad c_{j_k} = \langle b_j, S_{k-1}\rangle, \quad i_k = \operatorname*{argmax}_j |c_{j_k}|,$$

where $b_{i_k}$ is the receptive field of the neuron with index $i_k$ that was selected in step $k$ and $S_n$ is the residual after $n$ encoding steps. In each encoding step, the new residual $S_n$ is equal to the old residual $S_{n-1}$ minus the information that is encoded in the new base activation $c_{i_n}$. That is,

$$\begin{aligned} S_n &= S_{n-1} - c_{i_n} b_{i_n} \\ c_{i_n} b_{i_n} + S_n &= S_{n-1} \\ \langle b_{i_n}, S_{n-1}\rangle b_{i_n} + S_n &= S_{n-1}, \end{aligned}$$

where the terms on the left are orthogonal, since $b_{i_n}$ is orthogonal to $S_n$:

$$\begin{aligned} \langle b_{i_n}, S_n\rangle &= \langle b_{i_n}, S_{n-1} - c_{i_n} b_{i_n}\rangle \\ &= \langle b_{i_n}, S_{n-1}\rangle - \langle b_{i_n}, c_{i_n} b_{i_n}\rangle \\ &= c_{i_n} - c_{i_n}||b_{i_n}||^2 = 0 \end{aligned}$$

Note that the receptive fields $b_j$ have unit norm $||b_j|| = 1$. Then, the energy before and after encoding is conserved as:

$$||S_{n-1}||^2 = ||S_n||^2 + ||c_{i_n} b_{i_n}||^2 = ||S_n||^2 + |c_{i_n}|^2. \quad [3]$$

By applying Eq.3 recursively, we can decompose the energy of the retinal response $||R||^2 \equiv ||S_0||^2$ as:

$$||S_0||^2 = ||S_N||^2 + \sum_{k=1}^{N} |c_{i_k}|^2,$$

$$||R||^2 = ||S||^2 + ||C||^2. \quad [4]$$

This constitutes the energy conservation property of the matching pursuit algorithm that is employed to compute the reward for the vergence reinforcement learner (see Methods section).

**Accommodation Reward.** To better understand why maximizing the squared response of the retinal representation maximizes its entropy, we employ Plancherel's theorem:

$$||R||^2 = ||\tilde{x}||^2,$$

where $\tilde{x}$ is the Fourier transform of the retinal response. We can now rewrite $\tilde{x}$ as the successive convolution of the original image with a blur filter followed by the whitening filter. The blur filter can be thought of as an approximation of the point spread function due to image defocus, while the whitening filter can be associated with the processing of the retinal input by ganglion cells (see *Whitening* section):

$$||\tilde{x}||^2 = ||\tilde{W}\left(\tilde{G}\tilde{I}\right)||^2 = ||\tilde{G}(\tilde{W}\tilde{I})||^2.$$

Here, $\tilde{I}$ is the Fourier transform of the original image, $\tilde{G}$ is the Fourier transform of the Gaussian blur filter, and $\tilde{W}$ is the Fourier transform of the whitening filter. The last equality holds since the convolution is commutative so we can first apply the whitening operation of the retina followed by the blur filter due to optical defocus. In our model, we assumed the whitening filter to be independent of the level of input blur. This is motivated by the fact that the shape of retinal ganglion cell receptive fields does not change in response to visual deprivation (30–32).

It is further assumed that the retinal ganglion cell response whitens the input images, in that its frequency spectrum becomes flat for lower frequencies before it gets cut off at higher frequencies to minimize noise (14, 15) (compare *Whitening* section and Fig. S6). Then we can write

$$||\tilde{G}(\tilde{W}\tilde{I})||^2 \approx ||\tilde{G}c||^2 = c^2||\tilde{G}||^2,$$

where $c$ is a constant due to the whitening of the frequency spectrum. After reversing Plancherel's theorem we can now calculate the reward as:

$$\begin{aligned}
c^2||\tilde{G}||^2 = c^2||G||^2 &= c^2 \int_{-\infty}^{\infty} \left(\frac{1}{\sqrt{2\pi\sigma^2}} \exp\left(-\frac{x^2}{2\sigma^2}\right)\right)^2 \mathrm{d}x \\
&= \frac{c^2}{2\pi\sigma^2} \int_{-\infty}^{\infty} \exp\left(-\frac{x^2}{\sigma^2}\right) \mathrm{d}x \\
&= \frac{c^2}{2\pi\sigma^2} \sigma\sqrt{\pi} \\
&= \frac{c^2}{2\sqrt{\pi}} \frac{1}{\sigma}.
\end{aligned}$$

This demonstrates that the accommodation reward is inversely proportional to the standard deviation of the gaussian filter, i.e., decreases for increasing blur levels.

**Vergence Reward.** The vergence reinforcement learning agent aims to minimize the conditional entropy of the retinal representation given the information encoded at the cortical stage by maximizing it's negative: $-\mathcal{H}(R|C)$. Here, we show that by minimizing the patch averaged difference between the squared retinal response $||R||^2$ and the squared cortical response $||C||^2$ we maximize a lower bound of the conditional entropy.

We take the input to the sparse coding module as a sample of a multivariate random variable $R$. Then, for an estimator $\hat{R}$ the mean squared error is bounded from below by (compare (33), Ch. 8):

$$\begin{aligned}
E\left[||R - \hat{R}||^2\right] &\geq E\left[||R - E[R]\,||^2\right] \\
&= E\left[\sum_j^k (R_j - E[R_j])^2\right] \\
&= \sum_j^k E\left[(R_j - E[R_j])^2\right] \\
&= \sum_j^k \sigma_j^2,
\end{aligned}$$

where $||\cdot||$ is the Euclidean norm. The inequality follows from the fact that the mean of $R$ is the best estimator for $R$. Then, $\sigma_j^2$ is the response variance of the retinal ganglion cell with index $j$ and $k$ is the dimensionality of the system, i.e., the number of ganglion cells that encode the image.

Next, we introduce a multivariate Gaussian random variable $G$ with covariance matrix $\Sigma$ and entropy $\mathcal{H}(G)$. Further, its components $g_j$ are pairwise independent with variances $\sigma_j^2$. Then, the covariance matrix is diagonal and $\sigma_j^2$ are its eigenvalues.

It follows:

$$\sum_j^k \sigma_j^2 \geq \sum_j^k \ln\left(\sigma_j^2\right)$$
$$= \ln(\det(\Sigma))$$
$$= 2\mathcal{H}(G) - k\ln(2\pi e),$$

where the first inequality holds, since $x > log(x)\ \forall x > 0$. As the Gaussian distribution is the maximum entropy distribution for a given covariance matrix (33), it bounds the entropy of the retinal representation $\mathcal{H}(R)$, i.e.,

$$\mathcal{H}(G) \geq \mathcal{H}(R).$$

For an estimator with side information $Y$ it follows (33):

$$E\left[||R - \hat{R}(Y)\,||^2\right] \geq 2\mathcal{H}(R|Y) - k\ln(2\pi e). \qquad [5]$$

For our model, we define an estimator of the original input image as the reconstruction $\hat{R}_B(C)$ based on the cortical activities $C$ and the receptive fields $B$,

$$\hat{R}(Y) \equiv \hat{R}_B(C) = \sum_j c_j b_j,$$

where $b_j$ are receptive fields of cortical encoding neurons and $c_j$ is the respective activation after encoding. In principle, $\hat{R}_B(C)$ depends on both, the cortical activity vector $C$ and the matrix of receptive fields $B$, such that we would have to define $Y \equiv (C, B)$. However, we assume that the receptive fields adjust slowly compared to the neural activity. Then, the change in $\mathcal{H}(R|C, B)$ is dominated by the change of neural activity $C$ and we can simplify the left side of Eq. 5:

$$E\left[||R - \hat{R}_B(C)\,||^2\right] \equiv E[||S||^2] \qquad [6]$$
$$= E[||R||^2 - ||C||^2],$$

where we defined the residual image $S$ that is not represented at the cortical stage. The last equality holds due to the energy conservation property of the matching pursuit algorithm (see above), where $R$ is the retinal image before encoding and $C$ is the vector of cortical activities $c_j$. In every iteration, the vergence reward is the negative reconstruction error, i.e., $-||S||^2 = ||C||^2 - ||R||^2$, averaged over all $300 \times 2$ patches of the two scales. Therefore, over multiple iterations, an empirical estimate of the reconstruction error $E\left[||R - \hat{R}_B(C)\,||^2\right]$ is minimized, i.e., according to Eq. 5, we minimize an upper bound on a function that is monotonic in the conditional entropy $\mathcal{H}(R|C)$.

**Physiological Substrates for Reward Calculation.** In our model, the reward for the vergence learner is the negative difference between the retinal response and the average response of cortical neurons: $-(||R||^2 - ||C||^2) = ||C||^2 - ||R||^2$. The receptive fields of the cortical neurons in our model are similar to those of simple-cells that are found in the primary visual cortex (15, 34) (compare Fig. S1). The difference between cortical and retinal response can be interpreted as a prediction error, where the activity of the lateral geniculate nucleus in the thalamus is predicted by cortical activity. There are multiple suggestions, how such a prediction error could be computed in the cortical microcircuit (35–38). Yet, interactions between the thalamus and primary visual cortex may already be sufficient to compute the vergence reward. In the lateral geniculate nucleus, the minority of synapses stem from retinal axons, while there is an abundance of cortico-thalamic feedback connections integrated into a rich microcircuit (39–42). The thalamus projects to the primary sensory and higher-order cortices (43), from where activity is dispersed throughout the brain (44), including many areas that are thought to be involved in the generation of eye movements (45–47).

In summary, this suggests that the vergence reward could be computed as early as in the thalamus, only requiring feedback from the primary visual cortex.

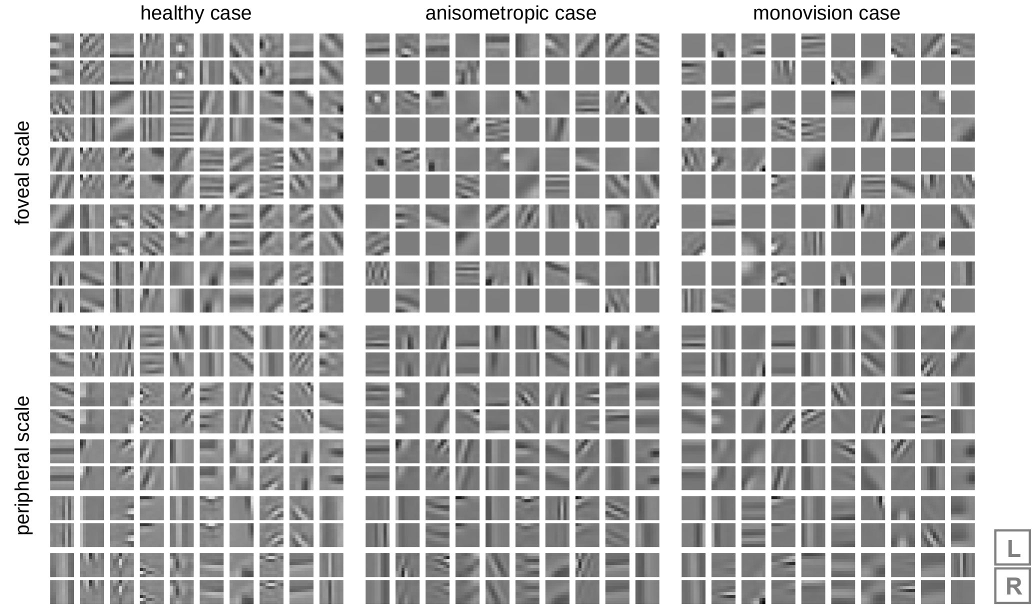


**Fig. S1.** $2 \times 8 \times 8$ px binocular receptive fields of cortical neurons after training for $5 \times 10^6$ iterations under different conditions (compare Main text, Fig. 3*C*). The statistics of the visual input was determined by the actions of the reinforcement learning agents, which were trained simultaneously. Receptive fields of the foveal scale are shown in the top row, receptive fields of the peripheral scale are shown in the bottom row. $50$ randomly selected neural receptive fields are shown per cortical encoder and condition. The upper patch represents the left eye's receptive subfields (L), the lower patch represents the right eye's receptive subfield (R). In the healthy case (left), binocular receptive fields evolve in both scales, with matching left and the right eye subfields. In the anisometropic case (center), in the foveal scale (center, top), receptive fields are mostly selective for input from the unimpaired, left eye while being unselective for input from the right eye (compare Main text, Fig. 5*C*, bottom). This is due to the sustained suppression of the right eye at the foveal scale (compare Fig. S9*A*). However, the peripheral scale is not suppressed (Fig. S9*A*) and binocular receptive fields emerge (center, bottom). This matches psychophysical findings in human patients (3, 4, 6, 49, 50) and receptive field measurements in animals raised with monocular deprivation (see (51) and references therein). In the monovision case, at the foveal scale, neurons are tuned towards one or the other eye (right, top), i.e., the receptive fields are either left or right monocular. This is similar to receptive fields of kittens which were raised under monocular deprivation (51, 52). In the peripheral scale, receptive fields tuned for lower frequencies are mostly binocular. Receptive fields tuned for higher frequencies are selective for one or the other eye, comparable to the foveal scale (right, bottom).

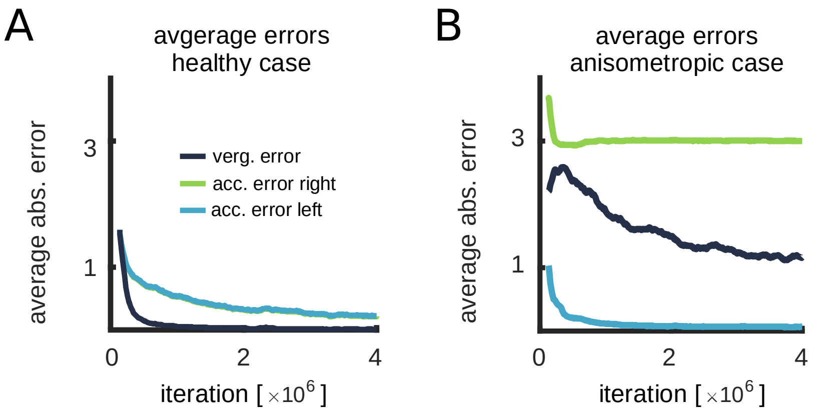


**Fig. S2.** *(A)* Running average of absolute (abs.) vergence and accommodation errors during healthy training measured in the final iteration before object change. The error gives the average distance between the respective planes and the object plane. The left and the right accommodation errors overlap since the left and the right accommodation planes coincide. The vergence error becomes almost zero. *(B)* Same as *A* for the anisometropic case. The model learns to focus the object with the left eye, leading to defocused input for the right eye. Vergence behavior is impaired compared to the healthy case in *A*.

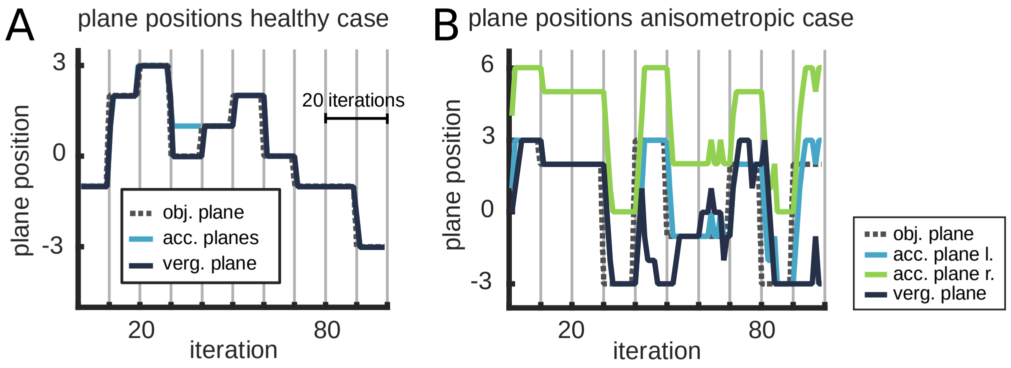


**Fig. S3.** Example trajectories of plane positions after training. Vertical gray lines indicate the end of fixations, where stimulus distance and image are randomly switched. *(A)* In the healthy conditions the left and right accommodation planes coincide (Main text, Fig. 3*C*, top). Shortly after the start of a new fixation (vertical gray line) all planes overlap at the object distance (dotted line) which results in sharp, zero disparity input for the left and right eye (compare Main text, Fig. 3*D*, top left). *(B)* Same as in *A* for anisometropic case (compare Main text, Fig. 3*C*, center). The object is continuously focused with the unimpaired (left) eye. Vergence performance is decrease compared to the healthy case. In this configuration, position $0$ is the closest distance that can still be focused by the right eye (compare Main text, Fig. 3*C*, center).

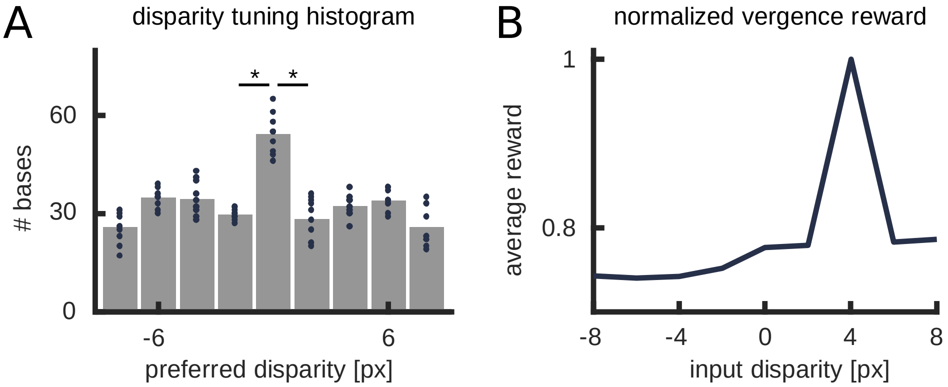


**Fig. S4.** *(A)* Disparity tuning of receptive fields from 10 dictionaries trained with uniform input disparity distribution for $100$k iterations. The preferred disparity of each receptive field was determined as the disparity that yields the largest average scalar product with a whitened input patch. Responses were averaged over 300 textures $\times$ 81 input patches. Each dot represents the number of receptive fields in a particular disparity bin of one cortical encoding dictionary. We tested if bin size distributions were significantly different between neighboring bins and marked them accordingly (*, p<0.0001, Kolmogorov–Smirnov test). This means that even though unbiased input with uniform disparity distribution was presented, receptive fields tuned to zero disparity are overrepresented. Therefore, the cortical encoding algorithm is not unbiased, i.e., $I(R, C)$ is not independent of disparity. Thus, when maximizing $-\mathcal{H}(R|C) = I(R, C) - \mathcal{H}(R)$, the sparse coder is not dependent on the lower entropy $\mathcal{H}(R)$ for zero disparity input. We can not exclude that this is due to the specific choice of our cortical coding algorithm, however, as long as there is no bias for another disparity that can offset the bias in $\mathcal{H}(R)$ the system should still favor zero disparity input. *(B)* We investigated the effect of introducing such a bias. We trained a set of bases with constant input disparity of $4$ px and measured the vergence reward for inputs with different disparities (averaged over $300$ textures). This resembles a scenario where early in life due to, e.g. mechanical constraints, the visual input is impeded. Receptive fields with non-zero disparity preference developed, which suggests that the system might develop strabismus. Closer analysis of this scenario is left for future work.

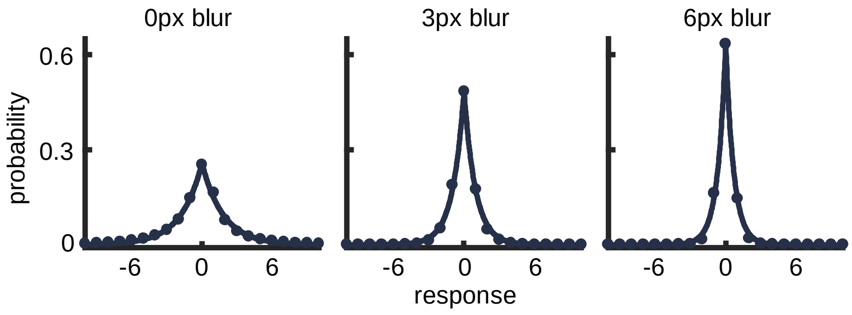


**Fig. S5.** *(A)* Estimated probability distribution of the entries of the whitened retinal response $R^*$ for different defocus blur values (blue dots). Average distribution over 300 images (bin size 1). Fitted with Laplace distributions (solid lines).

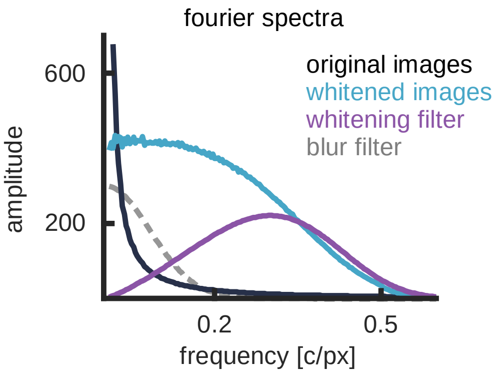


**Fig. S6.** Frequency spectra for whitened and non-whitened images, averaged over 300 textures, are shown in dark blue and light blue, respectively. The spectrum for the whitening filter is shown in purple. The $1/f^a$ decay of the non-whitened input image spectrum is balanced with the $f^a$ increase of the whitening filter, such that, after whitening, the spectrum becomes roughly flat for low frequencies. For comparison, the spectrum of a gaussian blur filter with a standard deviation of $2$ px is shown in dotted gray.

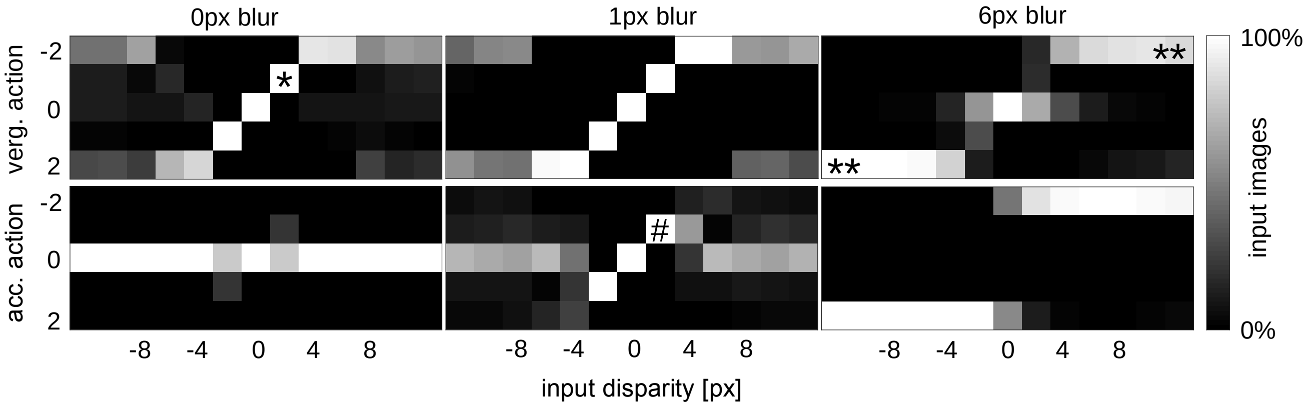


**Fig. S7.** Selected vergence (verg., top row) and accommodation (acc., bottom row) action for different input blur levels and disparities. The agent was trained under healthy conditions. Averaged histogram over 300 images. The intensity of each pixel indicates the fraction of input images with given blur and disparity for which the respective command was chosen. For zero blur and small absolute input disparities, the correct shift for the vergence plane is chosen. For example, for $+2$ px input disparity the vergence plane is shifted by $-1$, such that the input disparity would decrease by $2$ px and become zero in the next timestep (*, top left). Large input disparities are not resolved effectively if input blur is low. This is probably due to the low number of training samples, since low blur and high disparity become more and more unlikely the better the agent performs. For high blur levels and input disparities, the system can correctly reduce vergence error (**, top right). Since different blur levels give no defocus sign cue, the system utilizes disparity information to decrease the accommodation error. For example, in the case of $1$ px input blur and $+2$ px input disparity, the accommodation plane is moved by $-1$ (#, bottom center). Such a policy is successful in decreasing defocus blur if accommodation plane and vergence plane coincide. In fact, this is mostly the case after training (Fig. S3*A*).

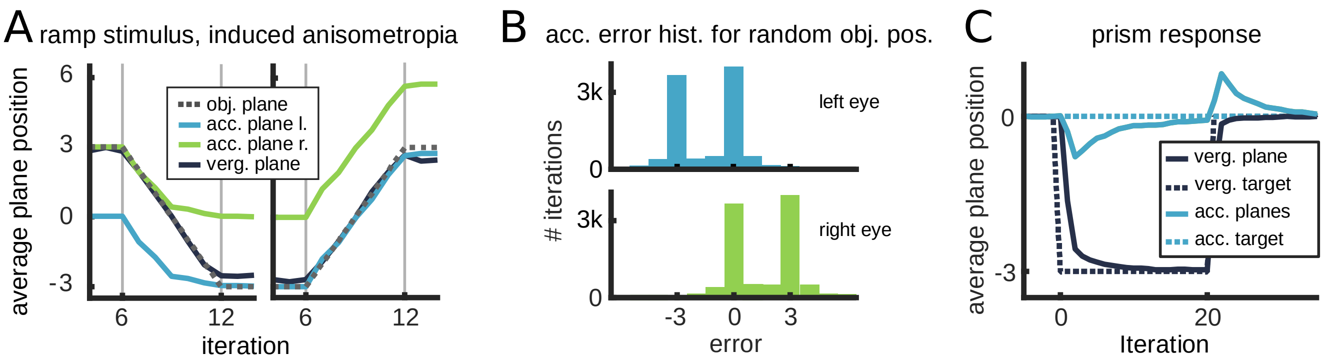


**Fig. S8.** Model responses to 'unnatural' input after training under healthy conditions. *(A)* Average plane trajectories for a ramp stimulus (dotted gray line) with induced anisometropia. By introducing a refractive error to the right eye, the left and the right accommodation plane are separated such that only one of the retinal images can be in focus at any given time (Main text, Fig. 3*C*, center). The stimulus position, i.e the object plane (dotted, gray line), gradually shifts from a distant position $3$ to a close position $-3$ (left panel) or vice versa (right panel). At each trial, the vergence and accommodation planes were initialized at fixed positions. The vergence plane was initialized such that it overlapped with the object plane. When the object shifted from a close (distant) to a distant (close) position, the accommodation plane of the left(right) eye was initialized to overlap with the object plane. As was previously suggested (53), the model chooses to focus with one or the other eye dependent on the initial stimulus position. When the object could no longer be focused with the right, myopic eye, it was focused by the left eye instead (left panel). *(B)* Accommodation error histogram of a model trained under healthy conditions and tested under lens-induced anisometropia for random object positions. The peaks at $-3, 0, 3$ indicate that most often, one or the other eye focuses the object. This results in an accommodation error of $0$ for the focusing eye and an absolute accommodation error that is equal to the anisometropic off-set due to the simulated lens for the other eye; In this case an absolute error of $3$ (Main text, Fig. 3*C*, center). There is a slight preference to use the unimpaired, left eye, to focus. This is probably because some distances at which the objects are presented are out of the right eye's focus range. Apart from that, both eyes are similarly recruited to encode the input. This indicates that despite induced anisometropia, due to the predominantly binocular receptive fields (Fig. S1), none of the eyes gets suppressed. *(C)* Average vergence and accommodation plane trajectories while adding and removing additional prisms at iteration $0$ and $20$, respectively. Due to the additional, virtual prisms in front of the eyes, the vergence target distance, which yields zero disparity (dotted, dark blue line) changed, while the accommodation target distance, which yields zero blur (dotted, light blue line), remained unaltered. In response, accommodation focus erroneously shifted in the direction of the new fixation target plane. When removing the prisms at iteration $20$, we observed the same effect in the opposite direction. This is in accordance with experiments (54) and can lead to visual fatigue and discomfort while using virtual reality headsets (55).

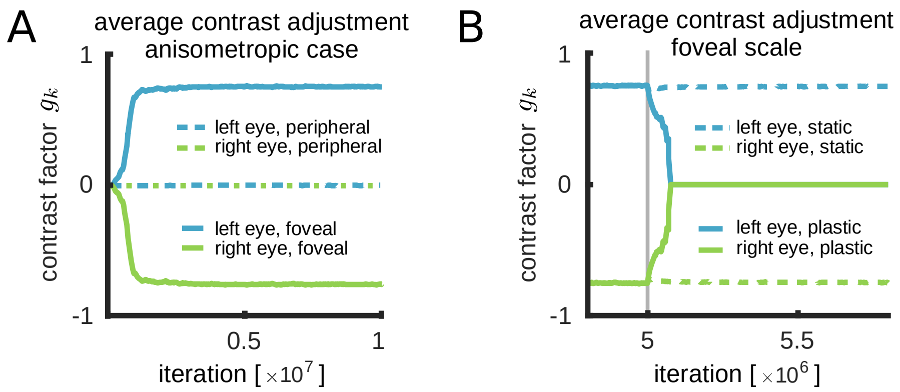


**Fig. S9.** Contrast adjustment factors $g_k = y_k - y_{\bar{k}},\ k \in \{l, r\}$ during anisometropic training and correction (compare Methods section). *(A)* Running averages of the contrast adjustment factors of peripheral (coarse) and foveal (fine) scale during training under the anisometropic condition. At first, suppression of the left and the right eye is balanced, i.e., there is no contrast adjustment for both eyes $g_l = g_r = 0$. Then, the left eye starts to dominate on the foveal scale. *(B)* Foveal scale contrast factors of formerly anisometropic model after correction of all refractive errors at iteration $5 \times 10^6$ (vertical gray line). The (dotted)solid line indicates a model with (non-)plastic receptive fields. The left (unimpaired) eye's contrast adjustment factor is shown in light blue and the right (hyperopic) eye's factor in green.

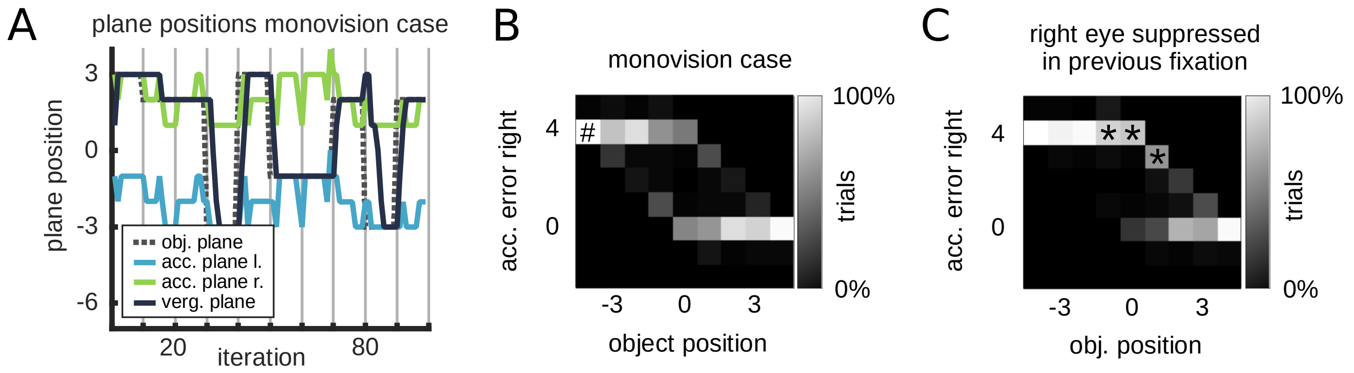


**Fig. S10.** *(A)* Example trajectories of plane movements at the end of training for the monovision case (compare Fig. S3*A* for healthy case trajectories and Fig. S3*B* for trajectories in the anisometropic case). The object plane is tracked by the vergence plane. When the object is positioned far away, it is tracked by the right (hyperopic) eye, when it is positioned close to the observer the left (myopic) eye is utilized. *(B)* Accommodation error of the right (hyperopic) eye at the end of a trial for different object positions in the monovision case with unbiased anisometropia. $n = 900$ trials with random object positions and image textures. In this configuration, position $0$ is the only distance that can be focused by both eyes (compare Main text, Fig. 3*C*, bottom). The error is defined as the difference between the right accommodation plane position and the object plane position. For close, i.e. negative object positions, the left (myopic) eye is favored which results in large accommodation errors for the right eye (#). At intermediate distances around zero, the history of the system, i.e. the current suppression, determines which eye is preferred (hysteresis): *(C)* Same as in *A* filtered for fixations where the right (hyperopic) eye was suppressed in the last iteration of the previous fixation, i.e., $y(x_l) > 0$ (compare *Suppression Mechanism* section). At intermediate distances, the non-suppressed left (myopic) eye is favored, which leads to a large accommodation error in the right eye (*).

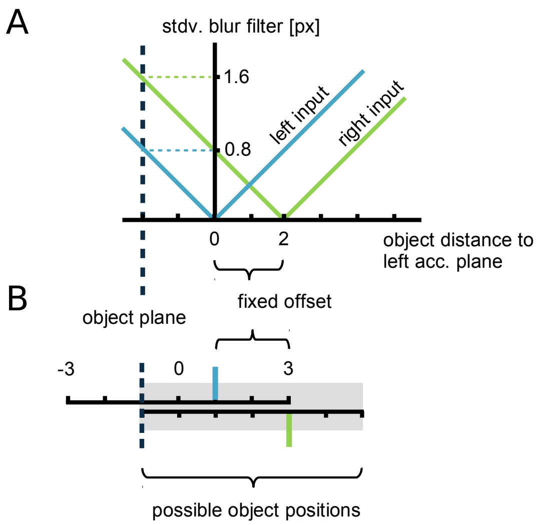


**Fig. S11.** *(A)* In our model, defocus blur is a function of the distance between the object plane and the two accommodation (acc.) planes. Defocus blur for the left retinal image is plotted in green and for the right retinal image in light blue and is realized with a Gaussian filter of variable standard deviation (stdv.). The dark blue dashed line indicates one exemplary object position. This configuration results in a large blur for the right eye input (dashed green line) and a lower blur level for the left eye (dashed blue line). In our model, defocus blur does not depend on whether the eye focuses in front of or behind the object. Therefore, there is no defocus sign cue in the healthy case where the left and the right accommodation plane coincide. Instead, input disparity is recruited to guide accommodation movements (see Fig. S7). In the anisometropic case, asymmetric blur levels for the left and the right eye provide a limited defocus sign cue which enables the agent to accommodate (compare Fig. S2*B*). For example, in Fig. S7*B*, the retinal image of the unimpaired (right) eye is much blurrier than the retinal image of the myopic (left) eye and the focal planes should be shifted towards the observer. However, note that this heuristic changes to the contrary when the object is positioned in between the two accommodation planes. *(B)* Plane configuration congruent with *A*. An object is placed at position $-1$ (dashed dark blue line). The left eye focuses at position $1$ (light blue line) while the right eye focuses at position $3$ (green line). Same scheme as in Main text, Fig. 3*B*.

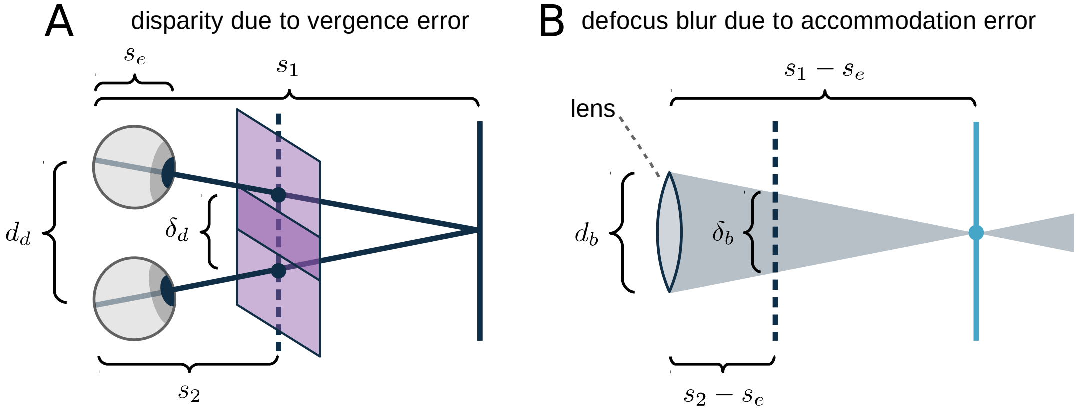


**Fig. S12.** Ratio between defocus blur and disparity. *(A)* An observer fixates at distance $s_1$ (dark blue line) while an object is presented at distance $s_2$ (dotted, dark blue line). $\delta_d$ is the distance between the two fixation points in the object plane (dark blue circles). The retinal input for the two eyes is indicated with two congruent squares (light purple). Then, $\delta_d = \frac{d_d}{s_1}(s_2 - s_1)$ is equal to the disparity shift between the left and right eye image. *(B)* Same configuration as in *B*, with the lens of one eye sitting at $s_e$, focusing at distance $s_2$ (light blue line). In this case, $\delta_b = \frac{d_b}{s_1 - s_e}(s_2 - s_1)$ gives the extent of space that is mapped to a single point on the retina and, therefore, determines the level of defocus blur. We now simplify and assume that the diameter of the eyeball $s_e$ is small compared to the object distance $s_1$. We further assume that $d_b$ is equal to the pupil size $p \approx 0.3$ cm (1). In the case of disparity, $d_d$ is approximately the interocular distance IOD $\approx 6$ cm (2). Then, the ratio between disparity and defocus blur is given as $\delta_d/\delta_b = d_d/d_b \approx \text{IOD}/p \approx 20.$

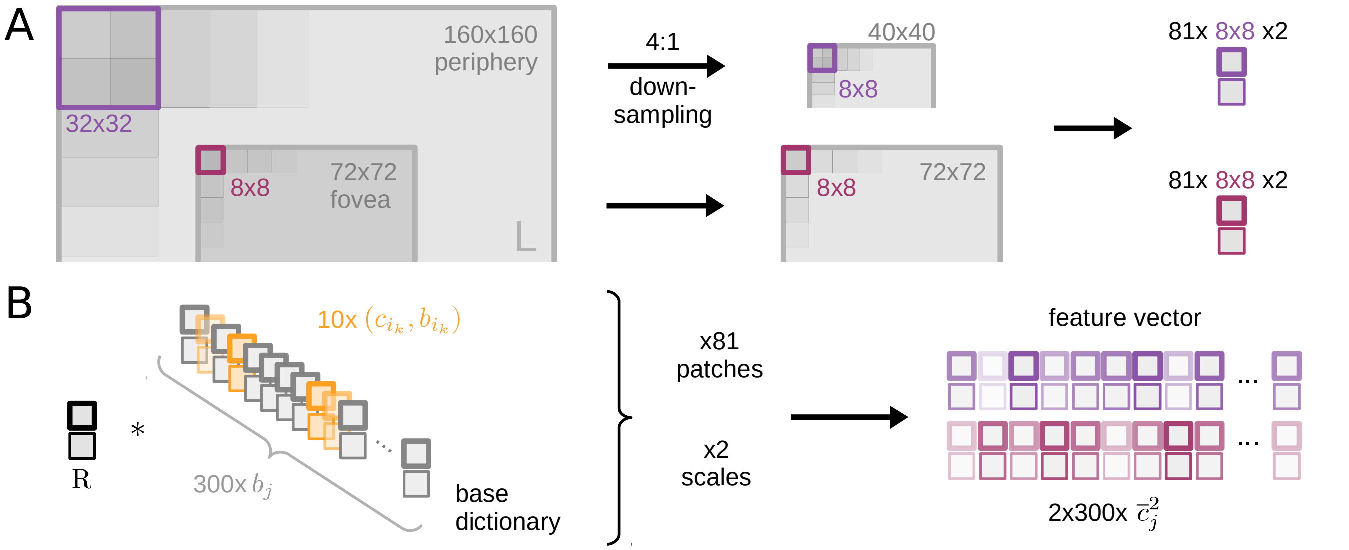


**Fig. S13.** Input processing *(A)* We extracted input patches at two scales. One peripheral scale (purple) and one foveal scale (red). For the peripheral(foveal) scale a $160 \times 160$ ($72 \times 72$) pixel-sized window was cut from the original image. The peripheral scale window was downsampled by a factor of $4$, using a Gaussian pyramid (48). Both windows were processed into $81$ binocular patches of size $8 \times 8 \times 2$ pixels. Due to the downsampling process, each peripheral scale patch represents an area of size $32 \times 32 \times 2$ pixels in the original window. Furthermore, each peripheral patch overlapped by half of its width with the neighboring patch. *(B)* Each scale had its own sparse coder with a separate dictionary. Every $8 \times 8 \times 2$ pixel binocular patch was normalized to zero mean intensity and subsequently encoded by 10 out of 300 bases $b_j$ using the matching pursuit algorithm (27). Thus, each one of the $162$ input patches was represented by the activities $c_{i_k}$ of $10$ bases of the respective dictionary. This resembles the sparse activity of simple cells in the visual cortex(15, 34). Selected receptive fields are marked in orange. Receptive fields of neurons with zero activity are shown in gray. Transparencies indicate activation strength. For each scale, the activity of the $300$ bases was squared and afterward averaged over all $81$ patches. The resulting activity vector can be understood as a model for complex cells in the visual cortex (23, 24). The two resulting $300$ dimensional feature vectors (right) were normalized to unit length, i.e., such that $\sum_j \bar{c}_j^2 = 1$, concatenated, and subsequently passed as the state information for the reinforcement learning agents (see Methods section).

### References


1. Spector RH (1990) The pupils in *Clinical Methods: The History, Physical, and Laboratory Examinations. 3rd edition.* (Butterworths).
2. Dodgson NA (2004) Variation and extrema of human interpupillary distance in *Stereoscopic Displays and Virtual Reality Systems XI.* (International Society for Optics and Photonics), Vol. 5291, pp. 36–46.
3. Bradley A, Freeman RD (1981) Contrast sensitivity in anisometropic amblyopia. *Investigative Ophthalmology and Visual Science* 21(3):467–476.
4. Holopigian K, Blake R, Greenwald MJ (1986) Selective losses in binocular vision in anisometropic amblyopes. *Vision research* 26(4):621–630.
5. Wang B, Ciuffreda KJ (2005) Blur discrimination of the human eye in the near retinal periphery. *Optometry and Vision Science* 82(1):52–58.
6. Babu RJ, Clavagnier S, Bobier WR, Thompson B, Hess RF (2017) Regional extent of peripheral suppression in amblyopia. *Investigative Ophthalmology and Visual Science* 58(4):2329–2340.
7. Van Essen DC, Newsome WT, Maunsell JH (1984) The visual field representation in striate cortex of the macaque monkey: asymmetries, anisotropies, and individual variability. *Vision research* 24(5):429–448.
8. Strasburger H, Rentschler I, Jüttner M (2011) Peripheral vision and pattern recognition: A review. *Journal of vision* 11(5):13–13.
9. Freeman J, Simoncelli EP (2011) Metamers of the ventral stream. *Nature neuroscience* 14(9):1195.
10. Priamikov A, Fronius M, Shi B, Triesch J (2016) Openeyesim: a biomechanical model for simulation of closed-loop visual perception. *Journal of vision* 16(15):25–25.
11. Klimmasch L, Lelais A, Lichtenstein A, Shi BE, Triesch J (2017) Learning of active binocular vision in a biomechanical model of the oculomotor system in *2017 Joint IEEE International Conference on Development and Learning and Epigenetic Robotics (ICDL-EpiRob).* (IEEE), pp. 21–26.
12. Lonini L, et al. (2013) Robust active binocular vision through intrinsically motivated learning. *Frontiers in neurorobotics* 7:20.
13. Teulière C, et al. (2015) Self-calibrating smooth pursuit through active efficient coding. *Robotics and Autonomous Systems* 71:3–12.
14. Atick JJ, Redlich AN (1992) What does the retina know about natural scenes? *Neural computation* 4(2):196–210.
15. Olshausen BA, Field DJ (1997) Sparse coding with an overcomplete basis set: A strategy employed by v1? *Vision research* 37(23):3311–3325.
16. Dayan P, Abbott LF (2005) *Theoretical Neuroscience: Computational and Mathematical Modeling of Neural Systems.* (The MIT Press).
17. Yoshioka T, Blasdel GG, Levitt JB, Lund JS (1996) Relation between patterns of intrinsic lateral connectivity, ocular dominance, and cytochrome oxidase-reactive regions in macaque monkey striate cortex. *Cerebral Cortex* 6(2):297–310.
18. Iacaruso MF, Gasler IT, Hofer SB (2017) Synaptic organization of visual space in primary visual cortex. *Nature* 547(7664):449–452.
19. Moradi F, Heeger DJ (2009) Inter-ocular contrast normalization in human visual cortex. *Journal of Vision* 9(3):13–13.
20. Carandini M, Heeger DJ (2012) Normalization as a canonical neural computation. *Nature Reviews Neuroscience* 13(1):51.
21. Said CP, Heeger DJ (2013) A model of binocular rivalry and cross-orientation suppression. *PLoS computational biology* 9(3):e1002991.
22. Brascamp J, Klink P, Levelt WJ (2015) The 'laws' of binocular rivalry: 50 years of levelt's propositions. *Vision research* 109:20–37.
23. Hubel DH, Wiesel TN (1962) Receptive fields, binocular interaction and functional architecture in the cat's visual cortex. *The Journal of physiology* 160(1):106–154.
24. Adelson EH, Bergen JR (1991) *The Plenoptic Function and the Elements of Early Vision*, ed. M. Landy and J. A. Movshon. (The MIT Press), pp. 3–20.
25. Wilson JR, Sherman S (1977) Differential effects of early monocular deprivation on binocular and monocular segments of cat striate cortex. *Journal of neurophysiology* 40(4):891–903.
26. Smith III EL, et al. (1997) Residual binocular interactions in the striate cortex of monkeys reared with abnormal binocular vision. *Journal of Neurophysiology* 78(3):1353–1362.
27. Mallat SG, Zhang Z (1993) Matching pursuits with time-frequency dictionaries. *IEEE Transactions on signal processing* 41(12):3397–3415.
28. Chandrapala TN, Shi BE (2014) The generative adaptive subspace self-organizing map in *2014 International Joint Conference on Neural Networks (IJCNN).* (IEEE), pp. 3790–3797.
29. Zhu Q, Triesch J, Shi BE (2017) Joint learning of binocularly driven saccades and vergence by active efficient coding. *Frontiers in neurorobotics* 11:58.
30. Sherman SM, Stone J (1973) Physiological normality of the retina in visually deprived cats. *Brain Research* 60(1):224–230.
31. Movshon JA, et al. (1987) Effects of early unilateral blur on the macaque's visual system. iii. physiological observations. *Journal of Neuroscience* 7(5):1340–1351.
32. Movshon JA, Van Sluyters RC (1981) Visual neural development. *Annual review of psychology* 32(1):477–522.
33. Cover TM, Thomas JA (2006) *Elements of information theory.* (John Wiley & Sons).

34. Hubel DH, Wiesel TN (1959) Receptive fields of single neurones in the cat's striate cortex. *The Journal of physiology* 148(3):574–591.
35. Rao RP, Ballard DH (1999) Predictive coding in the visual cortex: a functional interpretation of some extra-classical receptive-field effects. *Nature neuroscience* 2(1):79.
36. Spratling MW (2010) Predictive coding as a model of response properties in cortical area v1. *Journal of neuroscience* 30(9):3531–3543.
37. Bastos AM, et al. (2012) Canonical microcircuits for predictive coding. *Neuron* 76(4):695–711.
38. Sacramento J, Costa RP, Bengio Y, Senn W (2018) Dendritic cortical microcircuits approximate the backpropagation algorithm in *Advances in Neural Information Processing Systems.* pp. 8721–8732.
39. Erişir A, Van Horn SC, Sherman SM (1997) Relative numbers of cortical and brainstem inputs to the lateral geniculate nucleus. *Proceedings of the National Academy of Sciences* 94(4):1517–1520.
40. Alitto HJ, Usrey WM (2003) Corticothalamic feedback and sensory processing. *Current opinion in neurobiology* 13(4):440–445.
41. Wang W, Jones HE, Andolina IM, Salt TE, Sillito AM (2006) Functional alignment of feedback effects from visual cortex to thalamus. *Nature neuroscience* 9(10):1330.
42. Briggs F, Usrey WM (2011) Corticogeniculate feedback and visual processing in the primate. *The Journal of physiology* 589(1):33–40.
43. Schmid MC, et al. (2010) Blindsight depends on the lateral geniculate nucleus. *Nature* 466(7304):373.
44. Felleman DJ, Van DE (1991) Distributed hierarchical processing in the primate cerebral cortex. *Cerebral cortex (New York, NY: 1991)* 1(1):1–47.
45. Pierrot-Deseilligny C, Milea D, Müri RM (2004) Eye movement control by the cerebral cortex. *Current opinion in neurology* 17(1):17–25.
46. Hikosaka O (2009) *Basal Ganglia: Role in Eye Movements*, eds. Binder MD, Hirokawa N, Windhorst U. (Springer Berlin Heidelberg, Berlin, Heidelberg), pp. 350–352.
47. McDougal DH, Gamlin PD (2011) Autonomic control of the eye. *Comprehensive Physiology* 5(1):439–473.
48. Adelson EH, Anderson CH, Bergen JR, Burt PJ, Ogden JM (1984) Pyramid methods in image processing. *RCA engineer* 29(6):33–41.
49. Sireteanu R, Fronius M, Singer W (1981) Binocular interaction in the peripheral visual field of humans with strabismic and anisometropic amblyopia. *Vision research* 21(7):1065–1074.
50. Sireteanu R (1982) Human amblyopia: consequence of chronic interocular suppression. *Human neurobiology* 1(1):31–33.
51. Hunt JJ, Dayan P, Goodhill GJ (2013) Sparse Coding Can Predict Primary Visual Cortex Receptive Field Changes Induced by Abnormal Visual Input. *PLoS Computational Biology* 9(5).
52. Blakemore C (1976) The conditions required for the maintenance of binocularity in the kitten's visual cortex. *The Journal of Physiology* 261(2):423–444.
53. Bharadwaj SR, Candy TR (2011) The effect of lens-induced anisometropia on accommodation and vergence during human visual development. *Investigative Ophthalmology and Visual Science.*
54. Bharadwaj SR, Candy TR (2009) Accommodative and vergence responses to conflicting blur and disparity stimuli during development. *Journal of vision* 9(11):4.1–18.
55. Hoffman DM, Girshick AR, Akeley K, Banks MS (2008) Vergence-accommodation conflicts hinder visual performance and cause visual fatigue. *Journal of vision* 8(3):33–33.
